# Microbial Resuscitation and Growth Rates in Deep Permafrost: Lipid Stable Isotope Probing Results from the Permafrost Research Tunnel in Fox, Alaska

**DOI:** 10.1101/2025.01.20.633952

**Authors:** T. A. Caro, J. M. McFarlin, A. E. Maloney, S. D. Jech, A. J. Barker, T. A. Douglas, R. A. Barbato, S. H. Kopf

**Author notes:** Corresponding author: Tristan Caro.

## Abstract

Permafrost is at increasing risk of thaw as cold regions in the Northern Hemisphere continue to warm. Of particular concern is ice-rich, organic-rich, syngenetic “yedoma” type permafrost. The lability of organic carbon in permafrost post-thaw largely depends on the rate at which microorganisms resuscitate and proliferate after thousands of years in below-freezing, dark, anaerobic conditions. However, the resuscitation and growth rates of microorganisms in deep permafrost are unknown. To quantify these rates, we conducted lipid stable isotope probing (lipid-SIP) on permafrost cores collected from four locations within the Permafrost Tunnel near Fairbanks, Alaska. We compare rates of microbial growth, marker gene sequences, and greenhouse gas (CO_2_, CH_4_) emissions across cores held anaerobically at ambient and elevated temperatures. In deep, ancient permafrost, microbial biomass turnover is exceedingly slow, often undetectable, within the first month following thaw. Our results indicate microbial growth in response to anaerobic thaw has a notable lag period, where only 0.001 - 0.01% of cells turn over per day. This suggests a ‘slow reawakening’ that could provide some buffer between anomalous warmth and C degradation if permafrost refreezes seasonally. However, within six months, microbial communities undergo dramatic restructuring and succession, producing communities that are distinct from both the emplaced ancient and overlying surface communities. These results have critical implications for predictions of microbial biogeochemical contributions in a warming arctic, especially as thaw proceeds into deeper and more ancient permafrost horizons.

**Plain Language Summary:** Permafrost, earth material like soil, rock, or ice continually frozen for more than two years, contains more organic carbon than is currently in the atmosphere as CO_2_. As the Arctic warms and permafrost thaws, ancient microbes can reactivate, allowing the degradation of organic carbon that has accumulated in permafrost over millenia, resulting in the release of greenhouse gases. In this work, we measured the rates at which permafrost microorganisms resuscitate during thaw and related these growth rates to changes in microbial community composition and greenhouse gas emissions.

**Key Points:**

- Microbial growth is extremely slow within the first 30 days of thaw. Temperature may drive which taxa are active, but not growth rates.
- Subsurface microbes show a preference for glycolipids over phospholipids, suggesting a possible cryotolerance adaptation.
- Ancient, entrapped gases may be the primary source of emissions in early thaw stages.

## 1 Introduction

Permafrost, soil, rock, or ice that has been frozen for at least two consecutive years, comprises roughly one quarter of land area in the Earth’s northern hemisphere and stores a massive quantity of Earth’s carbon, estimated to be twice as large as the amount of CO_2_ currently in the atmosphere (Hugelius et al., 2014). The Arctic is warming at approximately twice the global rate (Stocker, 2014) and this threatens to release these buried stores of organic carbon as CO_2_ or CH_4_ driven by increased microbial degradation. In Alaska, the majority of permafrost exists to depths exceeding 10 m (Jorgenson et al., 2008) and underlies over 80% of Alaska’s surface area. Near Fairbanks, discontinuous permafrost reaches depths of up to 97 m (Jorgenson et al., 2008). In continuous permafrost regions on Alaska’s north slope, permafrost depth has been measured in excess of 660 m (Jorgenson et al., 2008). Therefore, a massive fraction of permafrost volume exists in deep subsurface horizons that are inaccessible to surface operations and difficult or impossible to assess with remote sensing. As anthropogenic landscape disturbances and temperature increase across the Arctic, permafrost thaw will presumably proceed further into deep subsurface horizons. Therefore, studies investigating the response of subsurface permafrost are urgently needed to understand its response to warming Arctic conditions.

Within permafrost lies a biosphere in suspended animation. Microorganisms emplaced in permafrost at time of deposition (freezing) persist at sub-zero temperatures through low-energy maintenance and repair (Price & Sowers, 2004). While buried and frozen, permafrost-hosted microorganisms may exhibit low levels of metabolic activity, likely as a result of trace liquid water available in interstitial spaces even at sub-freezing temperatures (Bakermans et al., 2003; Gilichinsky et al., 1993; E. M. Rivkina et al., 2000). Thaw remobilizes water, a previously limited resource, thereby liberating both microorganisms and their attendant organic matter. Numerous studies have examined the identity and genomic characteristics of microorganisms in permafrost, as well as how these communities shift during thaw (Barbato et al., 2022; Burkert et al., 2019; Mackelprang et al., 2017; Romanowicz & Kling, 2022; Waldrop et al., 2023), proposing mechanisms for how these communities experience ecological succession as a result of initial deposition (properties of the paleoenvironment), environmental filtering i.e., selection), and thaw characteristics (Doherty et al., 2020; Ernakovich et al., 2022; Waldrop et al., 2023). During thaw, microbial communities re-assemble after extreme environmental selection imposed by energy limitation and dark, freezing conditions. This thaw period is marked by microbial resuscitation, activity, and finally, proliferation.

Despite numerous studies investigating microbial identity, diversity, and functional capacity in permafrost, the rates at which microorganisms grow under both ambient and thaw conditions are unknown. Anabolic activity, expressed as biomass production, is a critical parameter to assess ecosystem functioning, habitability, and ecological dynamics (Foley et al., 2024). Measurements of microbial anabolic activity provide estimates for resuscitation time – the lag before microbial growth resumes. Additionally, growth rate is a mediator of microbial community assembly: knowing how quickly microorganisms are growing informs theories of microbial community succession. Thus, estimates of microbial growth rates, in both freezing and thawing conditions, are essential for a predictive understanding of microbial contributions to permafrost degradation and greenhouse gas emissions.

To conduct this study, we applied lipid stable isotope probing (lipid-SIP) on samples collected from the Permafrost Tunnel operated by the U.S. Army Cold Regions Research and Engineering Laboratory (CRREL). The Tunnel, located in Fox, AK (64.951256°N, 147.620731°W) is an ideal study site because it provides access to deep permafrost horizons, ranging in age from the modern to the late Pleistocene (∼40 kya), allowing study across permafrost of varying age (Barbato et al., 2022; Burkert et al., 2019; Douglas et al., 2011; Leewis et al., 2020; Waldrop et al., 2023) (**Fig. 1**). Phospholipid fatty acids (PLFAs) have been applied as a measure of living microbial biomass in environmental systems. PLFAs are an optimal analyte to assess microbial community size and function due to the fact that they rapidly degrade upon cell death and can therefore provide quantitative estimates of standing and produced microbial biomass. (Å. Frostegård et al., 2011; Zhang et al., 2019). Hydrogen stable isotope probing of PLFAs has been demonstrated to be a sensitive measure of microbial growth in a variety of settings including soil (Caro et al., 2023), clinical settings (Kopf et al., 2016), and marine sediments (Wegener et al., 2012). We additionally include analysis of glycolipid fatty acids (GLFAs) due to their hypothesized role in cryoprotection (Casillo et al., 2019; Corsaro et al., 2017).

**Fig. 1:**
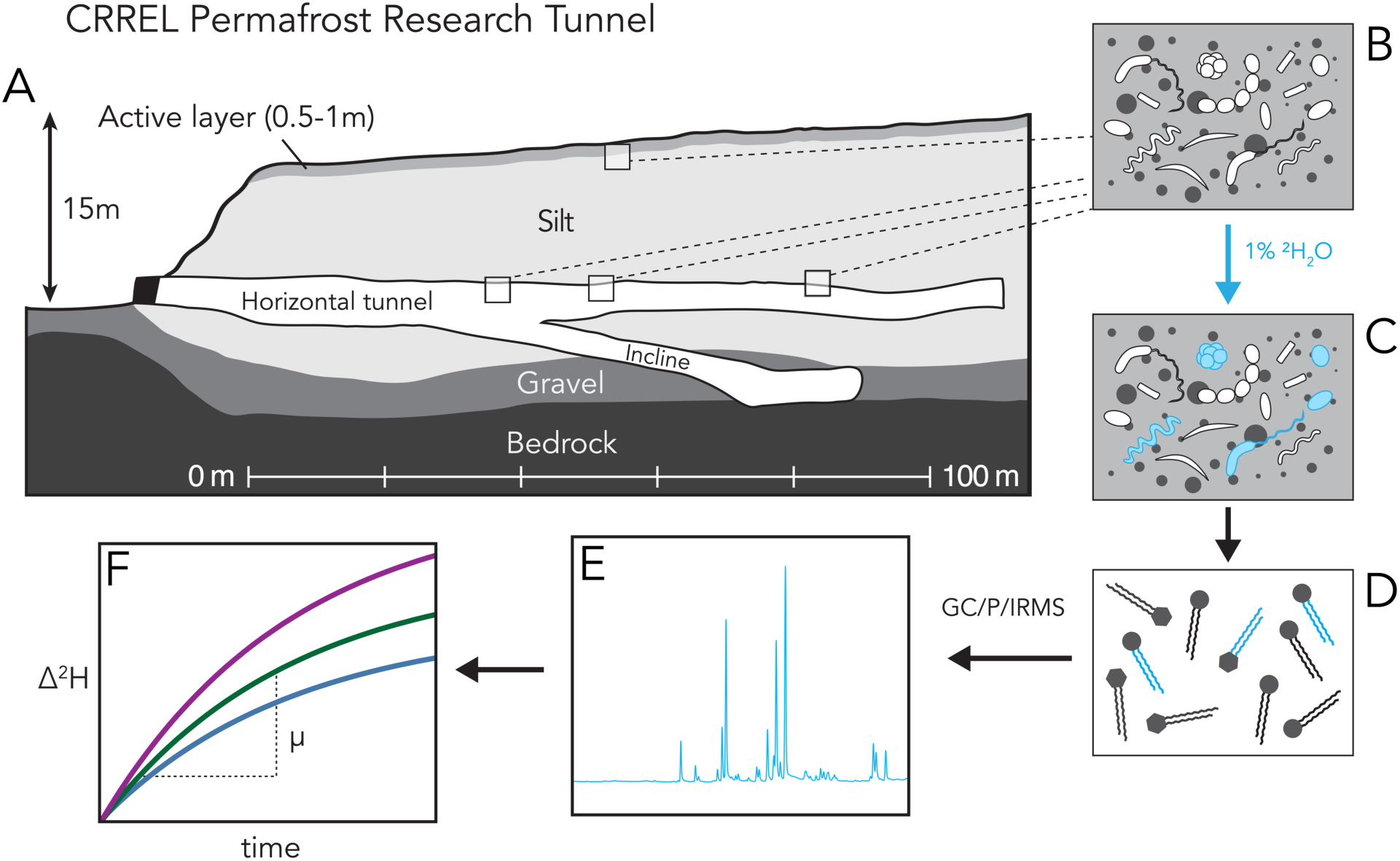
Schematic overview of study site and experimental procedure. (A) Cross section of the northern corridor of the CRREL Permafrost Tunnel in Fox, AK. The tunnel provides access to subsurface permafrost dated to the late Pleistocene. (B) Samples of permafrost from both surface and subsurface intervals contain relict microorganisms emplaced during initial permafrost development. (C) Lipid biosynthesis is a key expression of anabolic activity, and its detection represents microbial resuscitation and growth following thaw. Resuscitated microorganisms in the presence of deuterated “heavy” water tracer (1 at. % ^2^H) incorporate ^2^H into newly produced lipids. (D) Intact polar lipids are extracted from permafrost incubations. Lipids are separated into phospholipid (circle) and glycolipid (hexagon) fractions by headgroup polarity. (E) The deuterium composition of their fatty acid chains are analyzed by GC/P/IRMS. (F) The rate of deuterium enrichment relative to the beginning of the experiment (Δ^2^H) is used to calculate the production rate of microbial lipids (µ), either at the compound-specific or community-level scale.

To date, the absolute growth rates of permafrost microorganisms have never been measured. To address this key knowledge gap, we pursued a study of surface and subsurface permafrost with three objectives: (i) measure the biomass turnover of microorganisms at both ambient and thaw conditions, (ii) compare anabolic rates to 16S rRNA amplicon surveys to investigate the connection between microbial growth and community succession, (iii) evaluate the link between anabolic rates and the emission of greenhouse gases (CO2 and CH4).

## 2 Materials and Methods

### 2.1 Permafrost sampling and SIP incubation

Permafrost was collected from the U.S. Army Cold Regions Research and Engineering Laboratory Permafrost Tunnel located in Fox, Alaska, on August 6^th^, 2021 (**Fig. 1**). Cores of 1m in length were recovered using a gas-powered SIPRE (Snow, Ice, and Permafrost Research Establishment) corer (Jon’s Machine Shop, Fairbanks, Alaska, USA). Above the permafrost tunnel, the peat and soil layers were first removed with an ethanol-sterilized knife and trowel. Cores extracted from within the tunnel itself were drilled horizontally into the pore-ice cemented silt of the tunnel wall at distances of 35, 54, and 83 meters form the tunnel portal (**Fig. 1**). After recovery, all cores were enclosed in polyvinylchloride (PVC) tubing and stored inside the tunnel, which is held at −3°C.

All personnel wore Tyvek suits, personal N95 respirators, and nitrile gloves during core collection to reduce potential contamination. The SIPRE corer and all tools were cleaned with 70% isopropyl alcohol unless otherwise mentioned. Cores were stored in insulated boxes containing ice packs (Thermosafe U-Tek) to maintain temperature for shipping. All samples were transported to and stored at the Institute for Arctic and Alpine Research (INSTAAR) at the University of Colorado Boulder and stored at −20°C until processing. Temperature loggers (iButton) were included in each shipment to confirm the internal temperature of the shipments never exceeded 0°C during transport.

In the laboratory, permafrost cores were sectioned using a diamond-blade table-top tile saw. The saw was operated in a lab at room temperature (∼20°C) samples were rapidly transferred from and returned to an under-bench −20°C freezer. Before each sample, the saw blade and surface were cleaned with ultrapure water and 70% isopropyl alcohol. Then, the saw blade and metal table surface were cleaned by sequential triple rinsing with methanol (MeOH), dichloromethane (DCM), and hexane in a fume hood, in order to remove organic contaminants. The saw was kept clean of dust and debris between samples by covering its work surface with combusted aluminum foil. To process the cores for incubation, a cylindrical subsection of the intact core was cut, its exterior pared away, wrapped in combusted aluminum foil, and stored at −20°C until incubation. When beginning the incubation, the core subsection was unwrapped and cut into 12 subsections, one for each incubation time point and temperature condition. For each incubation, a subsection of core was weighed and added to a 250mL bottle along with 50 mL of filter-sterilized anaerobic ^2^H_2_O (1 at. % ^2^H) and the bottle was sealed with a butyl stopper. Incubation headspace was flushed with N_2_ for 30 minutes before being placed in a temperature-controlled freezer/refrigerator (Koolatron) held at a temperature of either −4°C, 4°C, or 12°C. Samples were incubated for either 0, 7, 30, or 180 days. At the end of the experimental period, microbial growth was arrested by transferring incubations to a −70°C freezer.

At the end of the SIP incubation period, and before freezing, permafrost was transferred to 50 mL centrifuge tubes and excess water was separated from permafrost pellets by centrifugation. Excess incubation water was filtered through a 0.2µm syringe filter before storage. Isotopic composition of the incubation water was measured to account for the isotopic contributions of water native to the permafrost. Both permafrost and excess water were frozen at −70°C. Frozen permafrost pellets were lyophilized for 48hr and stored at −70°C until lipid extraction.

### 2.2 Headspace Gas Measurements

Incubation headspace CO_2_ and CH_4_ were measured on a Silco SRI 8610C Gas Chromatograph Flame Ionization Detector (GC-FID) equipped with a methanizer. 0.5 mL of gas was anaerobically sampled from the incubations and injected using a gas-tight syringe. The GC-FID was equipped with a 2m, 2mm ID, ShinCarbonST 80/100 column and held at 200°C for each run. Gas concentrations were determined by comparison to a standard mix of 1% CO, CO_2_, CH_4_, which was periodically injected to account for instrument drift.

### 2.3 Lipid extraction

Intact polar lipids were extracted from the lyophilized permafrost pellets using a modified Bligh & Dyer lipid extraction (Bligh & Dyer, 1959; Quideau et al., 2016). In brief, 5.0g of freeze-dried soil was added to a solvent-cleaned PTFE centrifuge tube. 50µg of 23:0 phosphatidylcholine (23:0-PC) internal standard was added to each tube prior to extraction. Three extraction mixes, Mixes A, B, and C were applied. Mix A consisted of 1:2:0.8 (v:v:v) dichloromethane (DCM), methanol (MeOH), and phosphate buffer (8.7g/L K2HPO4, 3.5mL/L 6N HCl). Mix B consisted of 1:2:0.8 (v:v:v) DCM, MeOH, and trichloroacetic acid (TCA) buffer (250mL 10% TCA, 250 mL H2O, 8.8 g KOH). Mix C is a 1:5 (v:v) mixture of DCM and MeOH, respectively. 20mL of each extraction mix was applied to the lyophilized pellet in sequence. The mixture was vortexed for 20 seconds, sonicated in an ultrasonic bath for 10 minutes, then centrifuged at 3000 G for 10 minutes. After each centrifugation, the supernatant was carefully pipetted off and added to a collection vial. The sample was extracted with Mix A twice, Mix B twice, and Mix C once. Phase separation was induced by the addition of 25mL ultrapure water followed by 25mL DCM. Phases were allowed to separate for at least 2 hours, then the organic (bottom) phase was removed by pipetting. All samples were dried in a TurboVap LV at 35°C under a stream of N_2_ and stored at −20°C prior to downstream steps. With each batch of samples extracted, a contamination blank was processed simultaneously. No contamination was observed in process blanks. All glassware was combusted at 450°C for 8 hours prior to use. All PTFE vials were triple solvent rinsed with MeOH, DCM, and Hexane prior to use.

### 2.4 Intact polar lipid separation and derivatization

First, to remove residual extraction water and polar impurities, total lipid extract (TLE) was run through a sodium sulfate sand column (500mg 80:20 w/w Na_2_SO_4_:silica sand). The cleaned TLE eluent was then collected and purified using silica gel chromatography (Quideau et al., 2016) to focus isotopic analyses on intact polar lipids derived from intact cells. Combusted silica solid-phase extraction (SPE) columns containing 500mg SiO_2_ were conditioned with the addition of 5mL acetone, then two additions of 5mL dichloromethane (DCM). TLE was redissolved in 0.5mL DCM and transferred to the SPE column. Neutral lipids were eluted with the addition of 5mL of DCM. Glycolipids were eluted by the addition of 5 mL acetone. Phospholipids were eluted by the addition of 5mL MeOH. Each fraction was collected separately in combusted vials.

The glyco- and phospholipid extracts were derivatized to fatty acid methyl esters (FAMEs) via base-catalyzed transesterification using methanolic base (Griffiths et al., 2010; Rodríguez-Ruiz et al., 1998). Transesterification was initiated by adding 2mL hexane and 1mL 0.5 M NaOH in anhydrous methanol to dry lipid extracts. The reaction mixture proceeded for 10 minutes at room temperature before being quenched by the addition of 140µL of glacial acetic acid and 1mL water. The organic phase was extracted three times with 4mL hexane and dried under a stream of N_2_. A recovery standard of 10µg 21-phosphatidylcholine (21:0 PC) was added to each lipid fraction before derivatization to assess reaction yield.

### 2.5 FAME quantification and identification

A Thermo Trace 1310 Gas-Chromatograph equipped with a DB5-HT column (30 m x 0.250 mm, 0.25µm) coupled to a flame ionization detector (GC-FID) was used to quantify FAME concentrations based on peak area relative to the 23:0-PC extraction standard and a palmitic isobutyl ester (PAIBE) quantification standard, respectively. FAMEs were suspended in 100µL n-hexane and 1µL was injected using a split-splitless injector (SSL) run in splitless mode at 325°C; column flow was constant at 1.2mL/min. The GC ramped according to the following program: 80°C for 2 minutes, ramp at 20°C/min for 5 minutes (to 140°C), ramp at 5°C/min for 35 minutes (to 290°C). The FID was held at 350°C for the duration of the run. Major peaks were identified by retention time relative to a Bacterial Acid Methyl Ester (BAME) standard (Millipore-Sigma) and a 37 FAME standard (Supelco). Due to the uncertainty associated with identifying double bond position and stereochemistry for FAMEs containing multiple double bonds, unsaturated and poly-unsaturated compounds are identified only tentatively based on the position and retention time relative to the above standards. FAMEs are referred to using the nomenclature z-x:y, where x is the total number of carbons in the fatty acid skeleton and y is the number of double bonds and their position (if known), while z-is a prefix describing additional structural features of the compound such as branching, methylation, and cyclization. FAMEs were quantified using an external standard curve of PAIBE (**Fig. S1**).

### 2.6 FAME hydrogen isotope analysis

The hydrogen isotopic composition of FAMEs was quantified on a Thermo Scientific 253 Plus stable isotope ratio mass spectrometer coupled to a Trace 1310 GC via Isolink II pyrolysis/combustion interface (GC/P/IRMS). Chromatographic conditions were as such: 40°C for 2 minutes, 20°C/min to 180°C, 5°C/min to 265C, 30°C/min to 330°C, hold for 1.84 min, He carrier flow rate was 1.2ccm, the programmable temperature vaporization inlet temperature was ramped from 40°C to 400°C. Peaks were identified based on retention order and relative height based on coregistration with GC-FID chromatograms. Peak areas and isotopic values for mono- and di-unsaturated FAMEs were integrated to reduce uncertainty in downstream calculations at the expense of reduced compound specificity (e.g., all 16:1 isomers are reported simply as ’16:1’). Measured isotope ratios were corrected for scale compression, linearity, and memory effects using standards of both natural abundance and isotopically enriched fatty acid esters of known isotopic compositions ranging from −231.2 ‰ to + 3972 ‰ vs. VSMOW (Supplementary Text). Memory (peak-to-peak carryover) was corrected for using our previously defined framework (Caro et al., 2023). Hydrogen isotope calibrations were performed in R using the packages *isoreader* (v.1.4.1) and *isoprocessor* (v.0.6.11) available at github.com/isoverse. Calibrated isotope ratios were corrected for the H added during derivatization to FAMEs: the hydrogen isotope composition of methanol used for base-catalyzed transesterification was measured by derivatizing with phthalic acid with a known H isotopic composition (Arndt Schimmelmann, Indiana University) via acid catalysis. The resulting phthalic acid methyl ester (PAME) was analyzed by GC/P/IRMS and this correction was applied to all measured FAMEs.

### 2.6 Growth calculations

Calculation of biomass synthesis based on FAME enrichment is possible because the hydrocarbon chains of intact polar lipids consist only of C-H bonds that are nonexchangeable on biological timescales, unlike the readily exchangeable H bound to O, N, P, and S in lipid headgroups, proteins, and nucleic acids. The ^2^H tracer is therefore stably incorporated into newly synthesized fatty acid tails during anabolic activity. Biomass synthesis (growth or turnover) rates are calculated as described previously (Caro et al., 2023; Kopf et al., 2016). In brief, lipid biosynthesis reflects a combination of growth (*µ*) and repair, where *r* refers to the specific biosynthesis rate (day^-1^). Our measurements provides an upper bound for the specific growth rate *µ* and lower bound for the apparent generation time *T_G_* of the microbial producers of a given lipid:

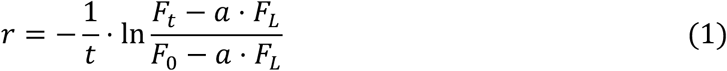

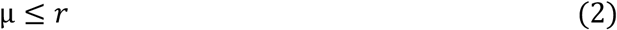

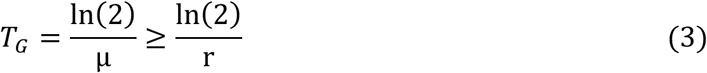

These calculations yield compound-specific growth/turnover rate and generation time estimates that can be viewed by themselves or aggregated into assemblage-level estimates of community growth by calculating an abundance-weighted mean:

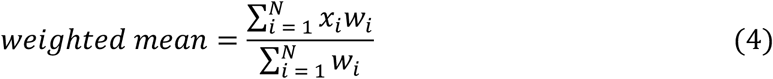

Where *x_i_* is the isotopic fractional abundance of compound *i* and *w_i_* is the relative abundance (weighting) of compound *i.* As discussed, these estimates aggregate cell-to-cell variations in growth. Uncertainty in each set of measurements was propagated through our calculations of *µ* by standard error propagation (Caro et al., 2023).

To assess microbial biomass production at extremely low rates, we took steps to increase our confidence in measurements of enriched ^2^H both at the compound-specific and assemblage (abundance weighted mean) levels. First, samples were followed by aggressive cleaning blanks where the instrument was ramped and held at a high (350°C) isothermal temperature followed by flushes of H_2_ reference gas. Second, isotopic compositions of individual FAMEs were corrected for memory effects as described in Caro et al., 2023, which reduces the possibility that ^2^H enrichment observed is the result of previous chromatographic peaks contaminating the peak of interest. Third, we define conservative thresholds for detection of deuterium enrichment and quantification of biomass turnover. We define our ^2^F detection limit as:

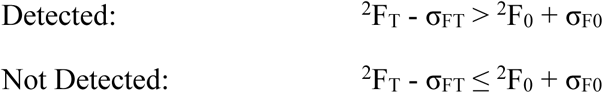

Where ^2^F_T_ is isotopic fractional abundance of a sample/compound at time of sampling (T), ^2^F_0_ is the isotopic fractional abundance of a sample/compound at the start of the experiment (T=0), and σ represents associated calibration error. For calculations of growth and turnover (*µ*), we apply standard error propagation. This propagated error results in an uncertainty term for biomass production rate (σ_µ_). We separately designate our limit of growth rate quantification as:

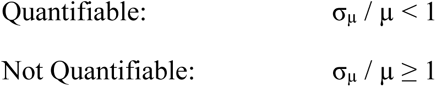

Where the fraction σ_µ_ / µ represents the relative error in the rate measurement. In other words, propagated error in growth rate must be less than the growth rate value itself to meet our criteria for quantification.

The terms ‘relative’ and ‘absolute’ growth rate are used according to Foley et al., 2024 where relative growth rate is the change in biomass or population relative to starting size (e.g., day^-1^) and absolute growth rate is the rate of change in mass per unit time (e.g., pg day^-1^). Relative growth rate measurements were converted to absolute growth rates by:

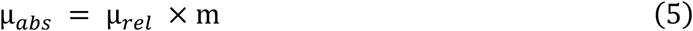

Where µ_abs_ is the absolute growth rate in units of mass per time per mass of permafrost [pg (day · g)^-1^ permafrost], µ_rel_ is growth rate in units of inverse time (day^-1^), and *m* is the permafrost weight-normalized lipid mass (pg lipid / g permafrost) measured by GC-FID and calculated based on the extraction and derivatization efficiency determined by internal standards (Materials and Methods Section 2.4). Cell abundances were estimated by multiplying PLFA and GLFA abundances by the mean conversion factor (3.5×10^-3^ pg PLFA/cell) estimated by A. Frostegård & Bååth, 1996. Resuscitated fraction (RF), the fraction of cells that resuscitate in a period of time, as:

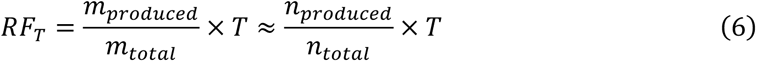

where m_produced_ is the mass of produced biomass within the time period of interest (T), m_total_ is the total biomass of the sample; n_produced_ and n_total_ are estimates of cell abundances converted from biomass. In this work, we report the fraction resuscitated after one day [RF_(1 day)_].

### 2.7 16S Ribosomal RNA Gene Sequencing

DNA was extracted from permafrost subsamples in triplicate and with negative controls (empty vials) using the DNEasy PowerSoil Pro DNA Isolation Kit (Qiagen) according to the manufacturer’s instructions with the following modifications: three triplicate extractions (via bead beating) were pooled onto a single elution column in order to increase the concentration of recovered DNA. DNA was cleaned (PCR inhibitors removed) with a DNEasy PowerClean Kit (Qiagen) with no modifications to the manufacturer’s protocol.

Extracted DNA samples were amplified in duplicate using Platinum II Hot-Start PCR Master Mix (ThermoFisher Scientific) SimpliAmp thermocyclers (ThermoFisher). The 16S rRNA gene primers 515F and 806R (V4 amplification) with Illumina sequencing adapters and unique 12-bp barcodes were used as described by the Earth Microbiome Project (Apprill et al., 2015; Parada et al., 2016). The PCR program was 94 °C for 2 min followed by 35 cycles of 94 °C (15 s), 60 °C (15 s), 68 °C (1 min), and a final extension at 72 °C for 10 min. Amplification was evaluated by agarose gel electrophoresis. Amplicons were cleaned and normalized with the SequalPrep Normalization Plate (Thermo Fisher Scientific) following the manufacturer’s instructions and then pooled together. Pooled libraries were quantified using both the Qubit ds DNA High Sensitivity Assay (ThermoFisher) and the KAPA Library Quantification Kit for Illumina platform (Roche). Sequencing was performed on an Illumina MiSeq using a v2 500 cycle PE reagent kit with paired-end reads at the Center for Microbial Exploration, University of Colorado Boulder.

To prepare samples for analysis with the DADA2 bioinformatic pipeline (Callahan et al., 2016), reads were demultiplexed with adapters and primers were removed using standard settings for cutadapt. We used standard filtering parameters with slight modifications for 2 x 150 bp chemistry where forward reads were not trimmed, and reverse reads were trimmed to 140 base pairs. A maximum error rate (maxEE) of 2 was allowed. Taxonomy was assigned with the SILVA (v138) reference database (Quast et al., 2013). Data was analyzed and visualized using the phyloseq and MicroViz packages in R (Barnett et al., 2021; McMurdie & Holmes, 2013).

### 2.8 Water stable hydrogen isotope analysis

The labeled incubation waters, following 0.2µm filtration through a PES membrane, were analyzed for the H stabe isotope composition (^2^F_L_) after gravimetric dilution with water of known isotopic composition (1:1000 w/w) to get into the analytical range of available in-house standards previously calibrated to Vienna Standard Mean Ocean Water (VSMOW) and Standard Light Antarctic Precipitation (SLAP). 1µL of each sample was measured on a dual inlet Thermo Delta Plus XL isotope ratio mass spectrometer connected to an H-Device for water reduction by chromium powder at 850°C (Putman et al., 2017). Measured isotope values in δ notation on the VSMOW-SLAP scale were converted to fractional abundances using the isotopic composition of VSMOW (R_VSMOW_ = ^2^H/^1^H = 0.00015576) (Meija et al., 2016) and corrected for the isotope dilution by mass-balance. The isotopic composition of each incubation was measured individually and resulted in values ranging between +30,065 – 49,153 ‰ vs. VSMOW (0.481 – 0.775 at. % ^2^H) These values were considered to be what cells encountered during tracer incubation (a combination of both the tracer and water present in the soil) and were applied to all growth rate calculations as *^2^F_L_*.

### 2.9 Permafrost dating and characterization

Approximately 5 g of lyophilized subsample of each permafrost core (prior to incubation with stable isotope probes) was sent for radiocarbon age dating at the National Ocean Sciences Accelerator Mass Spectrometry (NOSAMS) facility at Woods Hole Oceanographic Institution (Woods Hole, MA, USA), which uses NBS Oxalic Acid I (NIST-SRM-4990) as a reference standard. Collected data corresponds to NOSAMS accession numbers OS-175451, OS-175452, OS-175453, OS-175454. Routine geochemical description of the permafrost was conducted at the Colorado State University Soil, Water, and Plant testing facility (Denver, CO, USA). In brief, a KCl extract was used to quantify soil nitrate. An AB-DTPA extract was used to quantify P and S. Organic matter was determined by mass loss on ignition.

## 3 Results and Discussion

### 3.1 Experiment Summary and Site Characteristics

In **Table 1** we report key characteristics of the permafrost cores. Subsurface permafrost samples are of late Pleistocene age (37.9 – 42.4 kya). These samples exhibit low C:N ratios indicating substantial organic matter (OM; i.e., lignin) degradation and nitrogen accumulation. The surface permafrost (4.19 kya) sample exhibited a higher C:N ratio relative to the subsurface cores, which is consistent with less-progressed OM degradation in the surface setting (Mu et al., 2015). Because NO_3_^-^ was not detected in subsurface samples, total N likely represents ammonium accumulated as a result of microbial nitrate reduction.

**Table 1.**
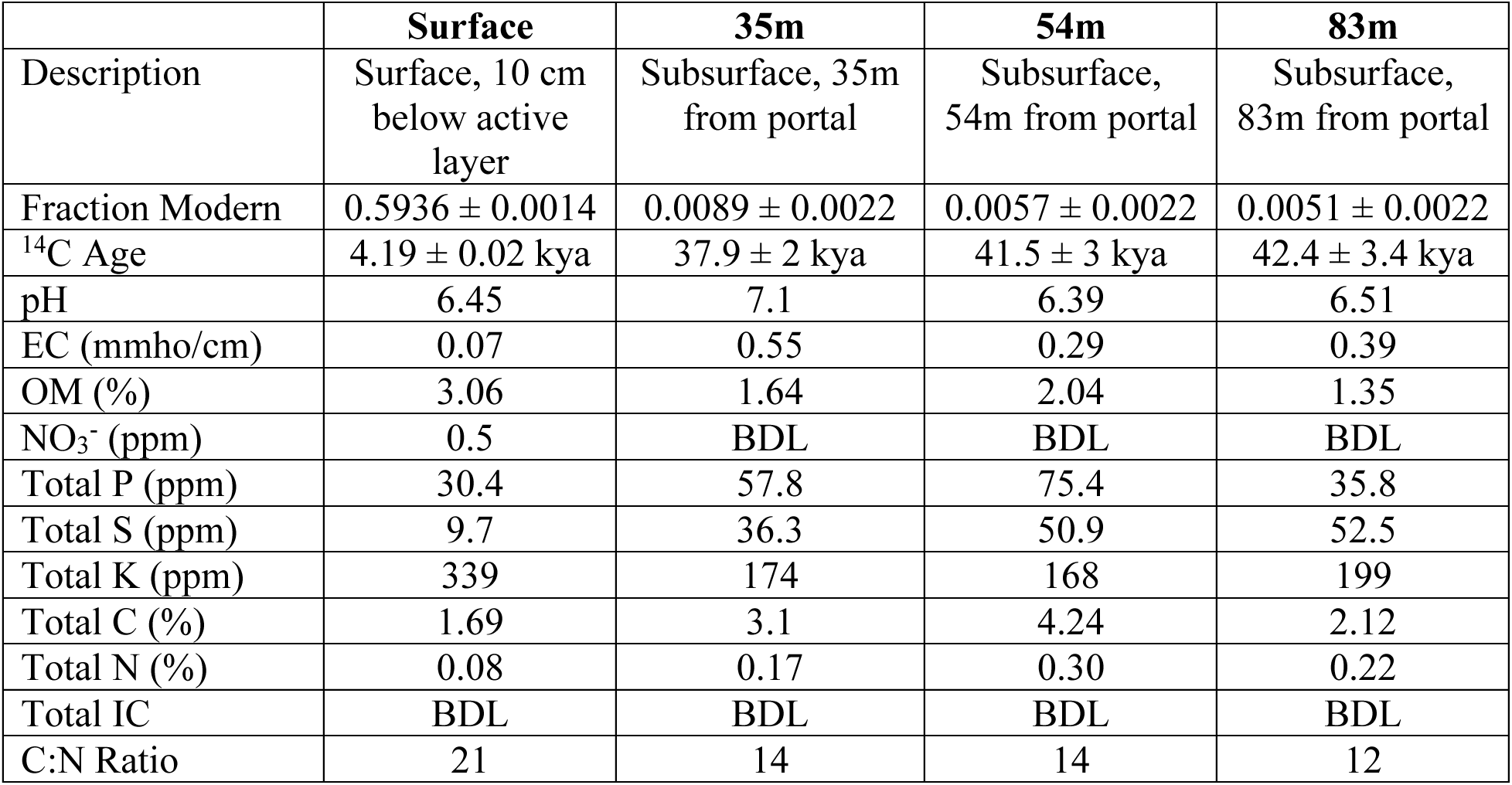
Summary of Site Characteristics. “BDL” indicates below detection limit.

|  | Surface | 35m | 54m | 83m |
| --- | --- | --- | --- | --- |
| Description | Surface, 10 cm below active layer | Subsurface, 35m from portal | Subsurface, 54m from portal | Subsurface, 83m from portal |
| Fraction Modern | $0.5936 \pm 0.0014$ | $0.0089 \pm 0.0022$ | $0.0057 \pm 0.0022$ | $0.0051 \pm 0.0022$ |
| $^{14}\text{C}$ Age | $4.19 \pm 0.02$ kya | $37.9 \pm 2$ kya | $41.5 \pm 3$ kya | $42.4 \pm 3.4$ kya |
| pH | 6.45 | 7.1 | 6.39 | 6.51 |
| EC (mmho/cm) | 0.07 | 0.55 | 0.29 | 0.39 |
| OM (%) | 3.06 | 1.64 | 2.04 | 1.35 |
| $\text{NO}_3^-$ (ppm) | 0.5 | BDL | BDL | BDL |
| Total P (ppm) | 30.4 | 57.8 | 75.4 | 35.8 |
| Total S (ppm) | 9.7 | 36.3 | 50.9 | 52.5 |
| Total K (ppm) | 339 | 174 | 168 | 199 |
| Total C (%) | 1.69 | 3.1 | 4.24 | 2.12 |
| Total N (%) | 0.08 | 0.17 | 0.30 | 0.22 |
| Total IC | BDL | BDL | BDL | BDL |
| C:N Ratio | 21 | 14 | 14 | 12 |

^14^C dates indicate that the subsurface samples collected overlap chronologically (Barbato et al., 2022; Burkert et al., 2019; Leewis et al., 2020). While the 35m, 54m, and 83m samples exhibit a positive trend with age, the higher error associated with carbon dating at these ages makes these values difficult to distinguish with confidence.

### 3.2 Microbial resuscitation and growth differ between surface and subsurface permafrost

In the surface permafrost collected here, microbial anabolic activity resumes fairly quickly after thaw (12°C) under anaerobic conditions, but the degree of activity is very low. ^2^H enrichment proceeds more rapidly among PLFAs compared to GLFAs in surface permafrost (in contrast with subsurface samples: see Section 3.4). The most rapidly produced PLFAs were identified as 16:1, followed by 16:0 and 18:1, whereas the most rapidly produced GLFAs were i-15:0, 16:0, and 14:0. Robust microbial growth can be observed after 30 days in both phospho- and glycolipid lipid pools from the surface core, and as early as 7 days in the phospholipid pool. ^2^H enrichments in PLFA pools correspond to abundance-weighted average growth rates between 2.1 × 10^-3^ after 7 days and 7.6 × 10^-3^ (day^-1^) after 30 days. The increase in growth rate observed (decrease in generation time) between the 7- and 30-day SIP incubations indicates a that microbial growth rate itself is increasing with time (**Fig. 2**). Growth rates correspond to community-level biomass generation times of 338 days (7 day incubation) and 91 days (30 day incubation). These generation times are substantially slower than those observed in alpine and tundra soil environments (14 – 45 days during a 7-day incubation) (Caro et al., 2023).

**Fig. 2:**
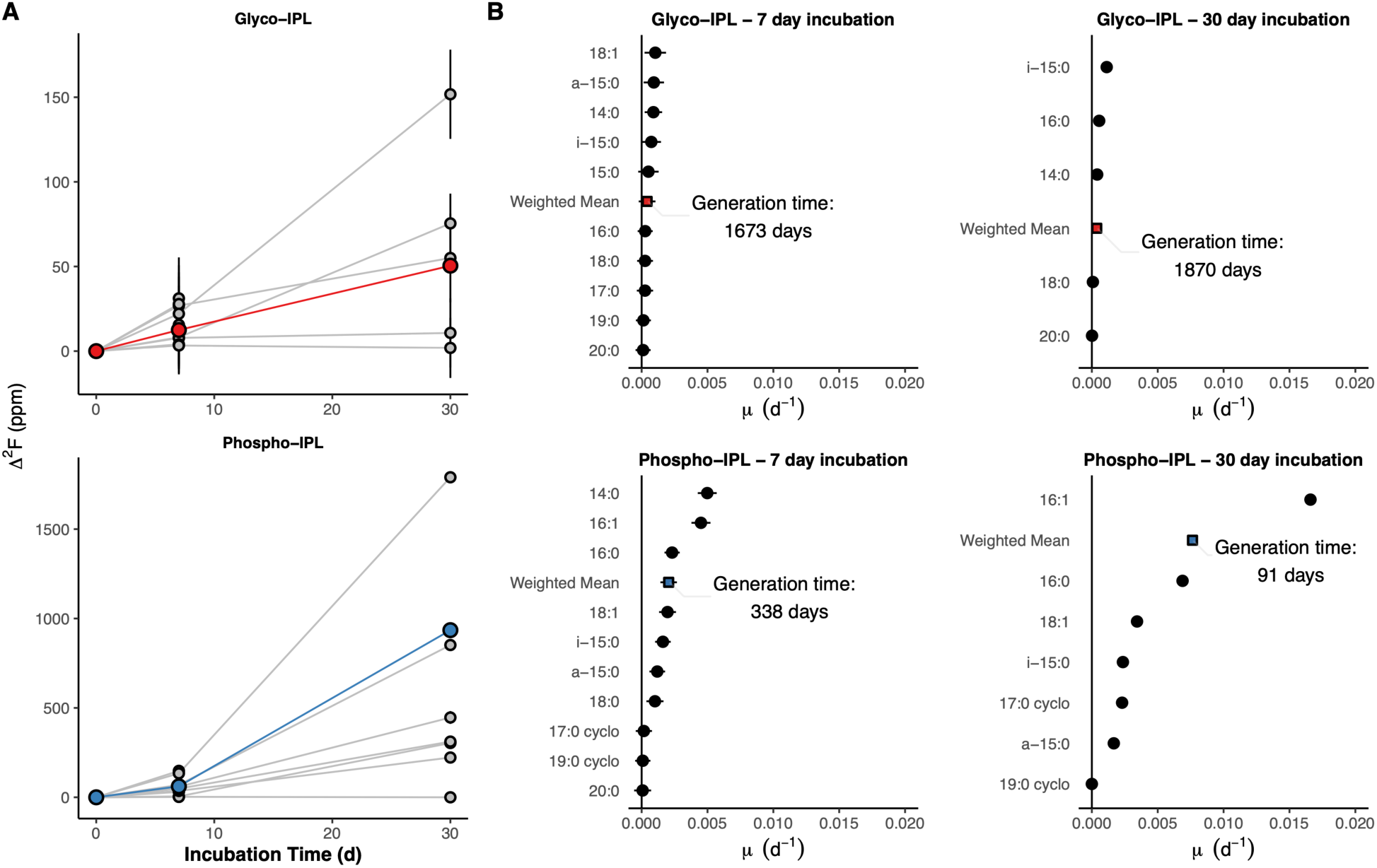
Compound-specific deuterium enrichment and inferred growth rates of microorganisms in surface permafrost thawing at 12°C. Panel A: compound specific deuterium enrichment (Δ^2^F) of lipid fatty acids over the course of the SIP experiment. Thin lines represent enrichment of individual lipid components. Bold lines in color represent abundance-weighted mean deuterium enrichment. Note different y axes for the two plots within panel A. Panel B: compound-specific growth rates of individual lipid fatty acids during both 7- and 30-day SIP incubations. Inferred generation time of the community-level weighted mean is indicated in each panel.

With reliable estimates of lipid biomass or cell abundances, measurements of relative growth rate (fractional replacement per unit time, e.g., day^-1^) can be converted into absolute measurements of growth rate (in units of biomass (g) produced per unit time). Because intact polar lipids are quantitative indicators of intact microbial biomass in the environment, we calculate absolute growth rate by multiplying the mass of lipids (PLFA or GLFA) recovered by their production rate. Measured PLFA content in surface samples varied between 3.8 – 7.1 × 10^3^ ng / g permafrost (Supplementary Table S3). Measurements of absolute growth rate in surface permafrost therefore range between 1.4 – 54 ng PLFA/day/g permafrost. Similarly, GLFA content in surface permafrost varied between 3.8 – 6.0 × 10^6^ pg GLFA/day/g permafrost, resulting in GLFA turnover rates of 1.4 – 2.5 ng GLFA/day/g permafrost.

In subsurface permafrost samples, microbial resuscitation and growth are much slower than surface samples and some are undetectable in the timescale of our 7 and 30 day incubations. For PLFAs, ^2^H enrichment is only detected in a small fraction of subsurface samples (**Table S2**). When detected, inferred relative growth rates lie between 3.3×10^-5^ to 5.7×10^-4^ (day^-1^), which correspond to community-level generation times of 3.3 – 57.5 years. The PLFA abundance of subsurface core samples was substantially lower than surface samples, between 330 to 1500 ng PLFA / g permafrost. Absolute rates of PLFA turnover in the subsurface therefore vary between 0 (undetectable) and 427 pg PLFA/day/g. Intriguingly, in subsurface samples, glycolipids exhibited more robust rates of turnover than phospholipids (68 – 624 pg/day/g permafrost), with all but 2 samples meeting detection criteria (**Fig. 3, Table S1**) (See Section 3.4). Subsurface samples exhibit no clear relationship between microbial growth, incubation temperature, and incubation duration (**Fig. 3**). In other words, some samples incubated at 12°C had lower rates of microbial growth than those incubated at 4°C and −4°C. Similarly, there were no clear trends between different aged subsurface permafrost and microbial growth (**Fig. 3**).

**Fig. 3:**
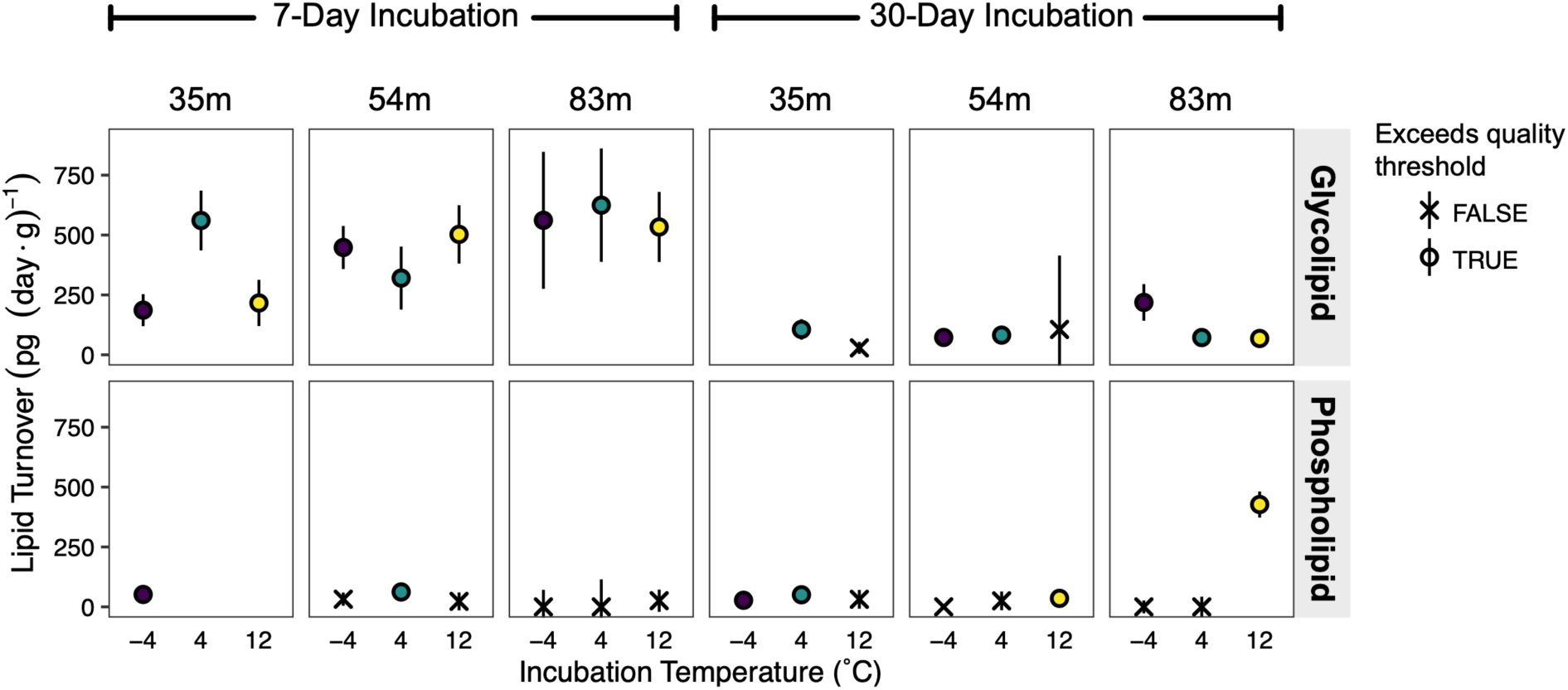
Microbial biomass growth in subsurface permafrost. Data symbols represent whether growth could be reliably quantified; data color corresponds to thaw temperature. Error bars represent total propagated error including instrument error, calibration error, and growth calculations. Missing data points indicate PLFAs/GLFAs were not detected. Data corresponding to the glycolipid and phospholipid fractions are plotted in the upper and lower panels, respectively. Each data point represents the weighted mean growth/turnover rate of all compounds detected in a given sample, normalized to the mass of permafrost in the incubation.

### 3.3 Quantifying the balance of dormancy and resuscitation

PLFAs and GLFAs provide quantitative estimates of living microbial biomass and cell numbers due to their rapid decay following cell death and demonstrated relationships between lipid content and cell concentrations in soil (A. Frostegård & Bååth, 1996; Zhang et al., 2019). Measurements of microbial growth via lipid-SIP can therefore be tied directly to absolute estimates of anabolic activity fractions across the microbial community, where the number of active cells out of the intact cell community can be estimated. For example, in the surface permafrost core, we convert PLFA content to estimates of intact cell abundance and estimate cell densities of 1.7-2.0×10^9^ cells/g permafrost. Based on our lipid-SIP dataset, we estimate that 4.0×10^5^ - 1.6×10^7^ cells are turned over daily over the course of 30 days.

If one assumes that microbial resuscitation is a discrete phenomenon (i.e., that measured isotopic growth is *only* the result of resuscitated cells), and that resuscitated cells contribute evenly to measured isotopic enrichment, then observed rates of biomass turnover can be a useful but imperfect metric of resuscitation rate. To this end, we calculate the resuscitated fraction (RF) of the microbial community with **Eq. 6** as the fraction of the microbiome that is resuscitated per day (RF_(1 day)_). In surface permafrost, the RF_(1 day)_ lies between 0.02 - 0.8 % of cells resuscitated per day. Resuscitation fractions in subsurface samples are substantially lower, ranging between RF_(1 day)_ = 0.0044 – 0.04 % as estimated by GLFA turnover and 0.0033 - 0.057 % by PLFA turnover. Exceptionally low RFs are evidence of high heterogeneity in resuscitation processes, suggesting that an equivalent of only 1 in 10,000 – 100,000 intact cells are successfully resuscitated per day in subsurface permafrost. The caveat with these estimates is that, when growth rates are aggregated into community-level (abundance-weighted) means, lipid-SIP cannot describe intracommunity heterogeneity i.e., the difference between one cell out of 100,000 undergoing complete turnover from 100,000 cells undergoing 10^-5^ turnover. However, we consider the calculation of RFs to be a useful and intuitive metric with which to frame microbial resuscitation and growth in this system. Indeed, in the anaerobic permafrost environment, high degrees of heterogeneity in resuscitation and growth is to be expected for dormant cells and spores, which use dormancy and non-uniform spontaneous resuscitation as a bet hedging strategy (Epstein, 2009; Lennon & Jones, 2011) and high resuscitation heterogeneity seems particularly likely given the observed trends in community composition (**Section 3.5, Fig. 4**).

**Figure 4:**
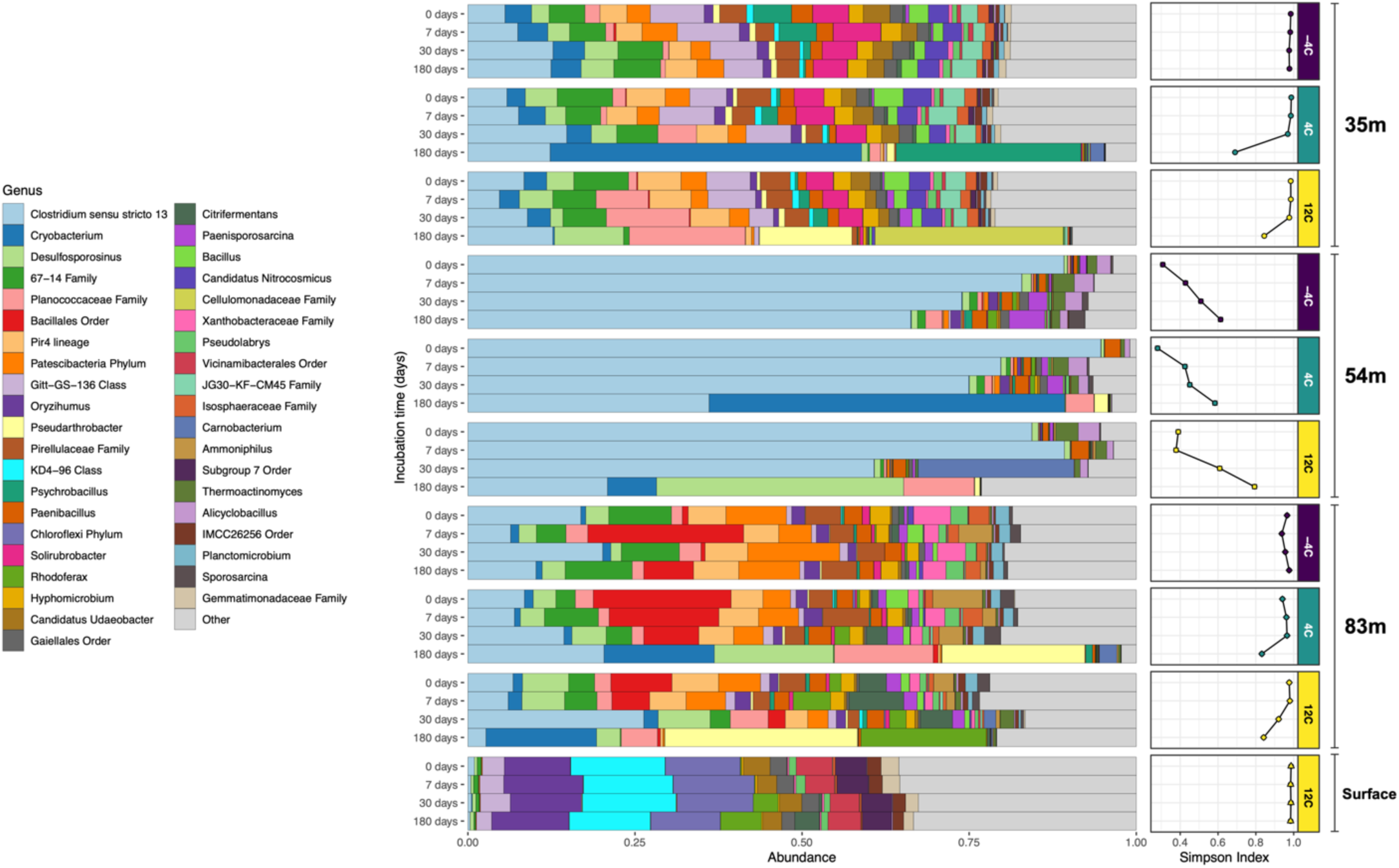
Microbial abundance across all samples and incubations. Horizontal axis of the main panel represents the fractional relative abundance of taxa grouped at the genus level (or highest annotated taxonomic rank, indicated in the key). Right-side panel indicates the Simpson alpha diversity index for each sample across incubations. Incubation time is noted on the y-axis; incubation temperature is noted in the right-hand panel labels.

While lipid-SIP has coarse taxonomic resolution, which is, at best, able to differentiate different phyla or broad functional groups (Caro et al., 2023; Willers et al., 2015b, 2015a), aggregating analytes from the entire microbial assemblage affords lipid-SIP the precision required to detect trace quantities of growth and estimate RFs. Single cell SIP analyses, such as those conducted with nanoscale secondary ion mass spectrometry (nanoSIMS) or Raman microspectroscopy (Caro et al., 2024; Eichorst et al., 2015), may not be suitable for permafrost environments due to the the potentially miniscule fraction of the microbial community that is resuscitated in the early stages of thaw.

### 3.4 Permafrost exhibits distinct patterns of microbial glycolipid and phospholipid production

There exists an intriguing distinction between production rates of PLFAs and GLFAs in the surface and subsurface permafrost. Compared to PLFAs, GLFAs are an order of magnitude more abundant across the subsurface samples (**Fig. 2**, **Fig. 3**) and produced at greater rates. GLFA production was detected in 88% of subsurface incubations, whereas PLFA production was detected in 38% of subsurface incubations (Supplementary Table S2). At the surface, the opposite trend is observed: GLFA production rates are far slower than their PLFA counterparts (**Fig. 2**), despite similar PLFA and GLFA concentrations (PLFA: 5 - 7×10^6^; GLFA: 4 - 6×10^6^, **Supplementary Table S1-S2**). We suggest that the abundance of glycolipids, and their more robust turnover in subsurface environments, points to the preference of cold-adapted bacteria to produce glycolipids as an adaptation to long-term freezing conditions. Indeed, glycolipids have been identified as membrane components of a variety of psychrophilic bacteria and are important constituents of lipopolysaccharide layers (LPS), which itself functions as a cryoprotectant (Casillo et al., 2019; Corsaro et al., 2017). It has been speculated that the high negative charge density of core LPS may be related to the sequestration of divalent cations, which may provide anti-freezing benefits by promoting the presence of liquid water at the ice-brine-cell interface (Corsaro et al., 2017; Gilichinsky et al., 1993). While we cannot unambiguously assign observed GLFAs to LPS structures (glycolipids may also serve as standalone cell membrane constituents), their robust production in comparison to phospholipids in subsurface cores supports the idea that glycolipids may serve a long-term cryoprotective role for organisms buried in permafrost.

### 3.5 In the long term, microbial communities undergo substantial restructuring after thaw

As described, exceptionally slow rates of microbial growth are observed after 7 and 30 days incubation post-thaw (Table S1, S2). SIP incubations of 180 days experienced dramatic changes in microbial community structure, lipid content, and lipid production. Lipids analyzed from the 180 day subsurface incubations were extremely enriched in ^2^H, up to 0.568 at.% (approx. δ^2^H_VSMOW_ = +35696 ‰). This observation is supported by striking increases in microbial biomass with time, across all incubation and temperature conditions (**Fig. S2 – S7**). Such isotopic enrichments approach that of the tracer solution, suggesting that prolific microbial growth has occurred, to the point where many constituents of the microbial community have undergone several doublings. The exact number of doublings cannot be estimated because isotope enrichment can no longer be quantitatively distinguished as cell biomass approaches isotopic parity with the tracer.

For the 35m and 83m cores, the alpha diversity (Simpson) index decreases over the duration of thaw, indicating the proliferation of specific taxa (**Fig. 4**). 180 days after thaw, in the 35m core, *Cryobacterium* and *Psychrobacillus* dominate at 4°C and *Desulfosporosinus, Planococcaceaea,* and *Pseudoarthrobacter* dominate at 12°C. Similarly, dominance of *Cryobacterium*, *Desulfosporosinus, and Pseudoarthrobacter* genera*; as well as Planococcaceaea* family, is observed after 180 days in the 83m core. The 54m core offers a striking contrast to the 35m and 83m cores (**Fig. 4**). This permafrost starts with an extremely homogenous microbial community dominated by *Clostridium sensu stricto 13*, a genus housing a variety of spore-forming species including *Clostridium bowmanii, Clostridium botulinum,* and *Clostridium huakuii.* Clostridia are more generally anaerobic spore-forming bacteria capable of fermenting soil organic matter (Boutard et al., 2014; Chades et al., 2018; Guan & Goldfine, 2021). Due to the dominance of a single taxa at the beginning of the incubation, the alpha diversity index (Simpson) increases over the course of thaw, as rare taxa begin to proliferate and balance the microbial community. After 180 days, the communities in core 54m see a substantial reduction in the original *Clostridium sensu stricto 13* community, with new abundance of *Cryobacterium* and *Desulfosporosinus.* Dominant taxa in these samples such as *Desulfosporosinus*, a genus of sulfate reducing bacteria, *Clostridium sensu stricto 13*, and *Psychrobacillus*, are widely described as spore forming (Buehler et al., 2018; Hippe & Stackebrandt, 2015; Krishnamurthi et al., 2010; Spring et al., 2003). The 16S data coupled with previously discussed resuscitation rates points to the relevance of heterogenous resuscitation (and the utility of RFs, as described above) because only a handful of community members, as low as 1 – 7 taxa, can emerge as dominant members of the microbial community.

The archaeal taxa detected in this study exhibit variation between cores and surface/subsurface (**Fig. S8**), but these communities do not exhibit clear successional trends with time. Archaea within the permafrost samples of this study comprise, at most, 4% of the microbial community and more typically ∼ 1% (**Fig. S8**). It is difficult to describe microbial succession of low-abundance microbial community members, as trace differences in abundance may be overprinted by the shifts of more abundant taxa. The 35m subsurface core is dominated by ammonium oxidizing archaea (AOA) *Candidatus Nitrocosmicus* Genus and Family *Nitrosphaeraceae.* Methanogenic archaea of genus *Methanobacterium* are the most abundant archaea in the 54m subsurface core as well as the surface core. The 83m core has low relative abundances of archaea but includes *Candidatus Nitrocosmicus* as well as Class *Bathyarchaeaia.* Bathyarchaeaia are notable as a diverse taxonomic group often associated with organic rich ecosystems and have been found capable of fermentation and acetogenesis of complex organic matter (Evans et al., 2015; He et al., 2016; Lazar et al., 2016).

Subsurface microbial communities become more similar over the course of thaw (**Fig. 5**). However, these communities do not converge towards a composition resembling that of younger permafrost, but rather, they form a “revenant” microbial community that is clearly distinct from modern surface permafrost-hosted microbial communities (**Fig. 5**). It is also important to note that the surface-hosted permafrost community maintained its structure over the course of thaw at 12°C and did not display indications of significant succession. These results suggest that surface permafrost sites may be poor proxies for predicting microbial community dynamics and responses in deeper horizons, which make up the bulk share of permafrost by volume (Jorgenson et al., 2008). The revenant community that we describe is the result of millennia of environmental filtering which includes the ancient depositional environment, duration of burial, metabolite, and energy limitation, etc. (Ernakovich et al., 2022; Waldrop et al., 2023). The convergence of subsurface microbial communities towards a shared revenant community suggests that common selection forces may be shaping the trajectory of communities during thaw. Further investigation of microbial succession dynamics in permafrost is required for a predictive understanding of community responses to thaw and the identification of taxonomic ‘indicators’ of thaw processes (Ernakovich et al., 2022).

**Figure 5.**
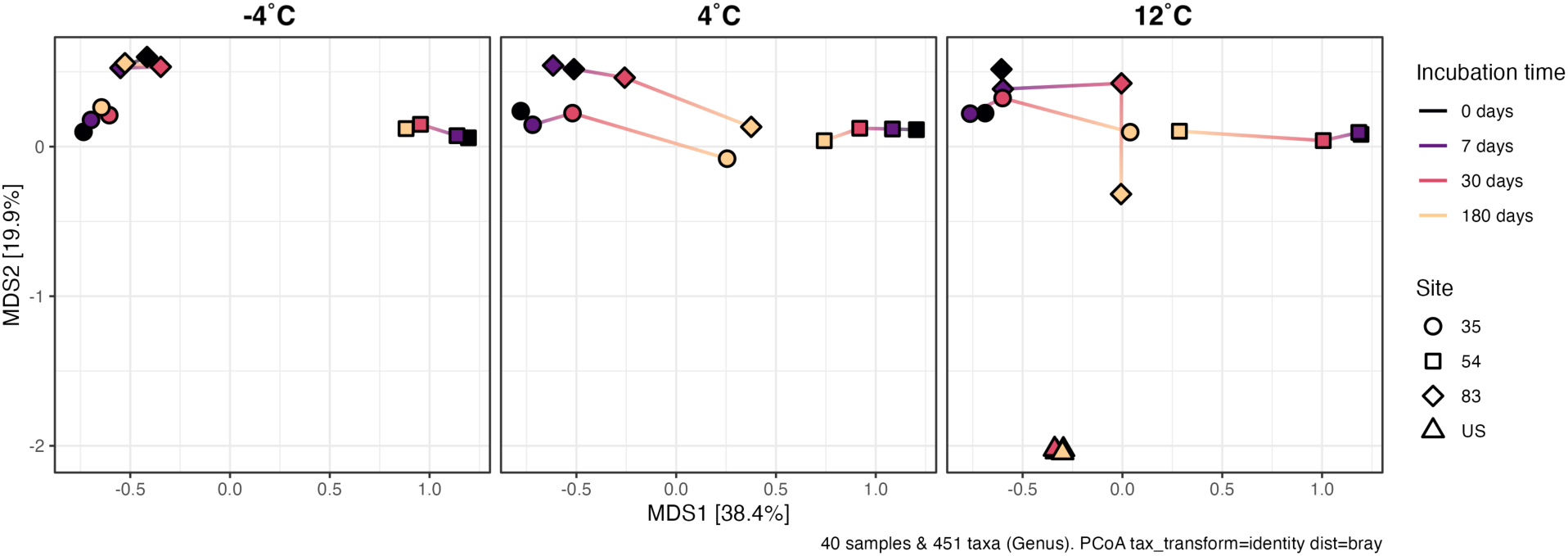
Microbial community similarity across incubation time series. Points are plotted based on Bray-Curtis dissimilarity. Point shape is the sampling site, where 35, 54, and 83 are distance from portal entrance in meters and “US” indicates Undisturbed Surface permafrost.

Our 16S amplicon survey, coupled with our lipid-SIP results, indicate that rates of subsurface microbial resuscitation and biomass growth do not relate to thaw temperature. It is also clear that subsurface microbial communities held at above-freezing temperatures under anaerobic conditions undergo dramatic community turnover and succession after 180 days, whereas those held below freezing do not (**Fig. 4**). The trace biomass turnover we detect at sub-freezing temperatures is not of sufficient magnitude to result in substantial community shifts (**Fig. 3**, **Fig. 4**) but supports previous observations of trace microbial activity at temperatures below freezing (Bakermans et al., 2003; E. M. Rivkina et al., 2000). Temperature does not appear to drive differences in growth rate, but does exert control over which taxa grow. Permafrost microbial communities host organisms with a wide range of growth temperature preferences, and the speed of this growth is likely driven by the difference between optimal growth temperature and thaw temperature, as has been suggested through *in vitro* studies of permafrost isolates, rather than temperature itself (Ernakovich & Wallenstein, 2015; Panikov & Sizova, 2007). This concept is suppoted by recent SIP studies of soil warming, which similarly support the finding that temperature affects the identity of growing taxa more than their growth rates (Metze et al., 2024).

### 3.6 Gas production is heterogenous and difficult to predict

Over the course of our anaerobic lipid-SIP incubations, we tracked the evolution of CO_2_ and CH_4_ in incubation headspace by GC-FID (**Fig. 6**). CO_2_ production was most rapid in surface permafrost, with a 30-day production rate of 3.1 × 10^-1^ nmol ml^-1^ g^-1^ day^-1^. CH4 emission was relatively muted in surface permafrost (2.1 × 10^-4^ nmol ml^-1^ g^-1^ day^-1^). In contrast, CH_4_ efflux was greater in subsurface samples (3.8 × 10^-4^ to 1 × 10^-2^ nmol ml^-1^ g^-1^ day^-1^). CH_4_ emission decreased with distance from the tunnel portal, but due to the similarities in permafrost age (**Table 1**) we are hesitant to attribute differences in methane emission to permafrost age or to microbial community composition (**Fig. S8**).

**Fig. 6.**
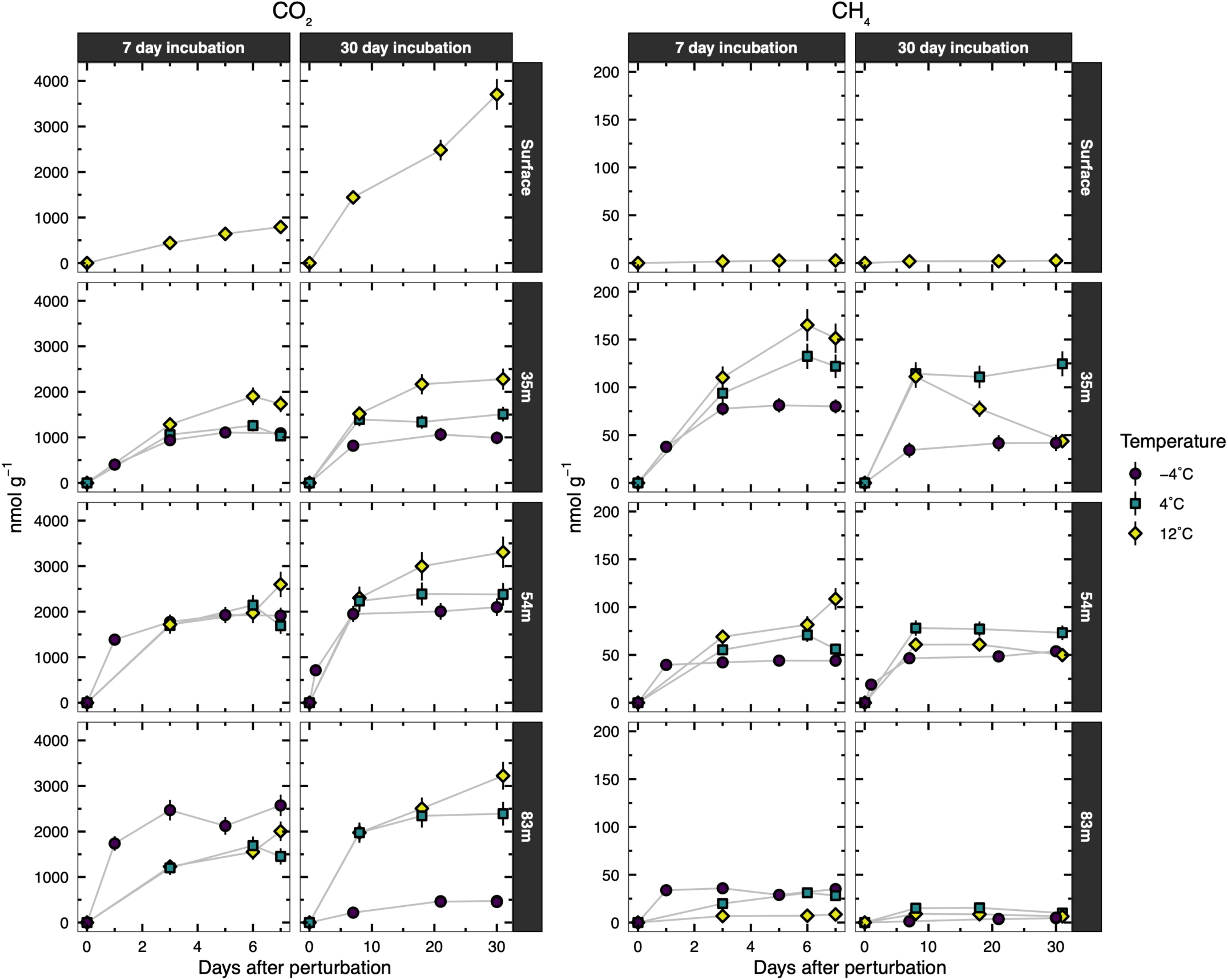
Carbon dioxide and methane efflux. Gas efflux is reported in nmol gas per gram of permafrost (wet weight). Error bars represent the standard error of the calibration. Note differences in horizontal axes between 7 and 30 day incubations as well as differences in vertical axes between CO_2_ and CH_4_ panels.

CO_2_ and CH_4_ production observed over the course of these incubations represents not only gases produced by biological activity during the course of the SIP incubation but also biogenic gas entrained within the permafrost matrix, accumulated over millennia and/or emplaced during initial deposition (E. Rivkina et al., 2007), liberated by the processes of thaw or the introduction of the isotopic tracer. These sources are difficult to disentangle. Our observed heterogeneity in gas production is likely the compounding of multiple factors: quantities of trapped gas, microbial community composition, thaw temperature, and availability of organic substrates and e^-^ donors/acceptors. In our study, the detection of methanogenic taxa by 16S amplicon sequencing does not clearly relate to methane emission (**Fig. 6**, **Fig. S8**). Indeed, it is possible that biogenic methane was not produced over the course of our incubation at all, and emissions here are instead from entrained methane, as methanogenic archaea may require substantial “priming” of their environment by bacteria to produce their necessary substrates regardless of anaerobic conditions (Ellenbogen et al., 2023). Previous anaerobic permafrost incubations have observed lag times of up to a year before methanogenic conditions are achieved (Yang et al., 2021). Heightened short-term methane emission observed in subsurface permafrost is therefore likely the result of biogenic methane accumulation over millenia.

### 3.7 Areas for future investigation

A limitation of this study is that microcosm incubations, even under tightly controlled analog conditions, cannot perfectly replicate the stochasticity of *in situ* permafrost thaw at the landscape scale. In a true thaw scenario, it is conceivable that increased hydrological connectivity would lead to increased rates of microbial dispersal. If dispersal is limited, environmental filtering may dominate the composition of the microbial community, but increased dispersal rates could lead to increased microbial community functioning due to the arrival of surface bacteria that are well-adapted to thawed conditions and potentially more capable of rapid growth – such organisms could quickly overprint an ancient community. As closed systems, our incubations do not experience thaw-induced input from surface microbial communities or hydrologic communication with surrounding subsurface permafrost environments. Furthermore, hydrological intrusion could supply sufficient oxygen that, in turn, could allow more rapid degradation of recalcitrant organic matter that is difficult or impossible to degrade anaerobically (Wilmoth et al., 2021). The smaller organic moieties produced may provide a suitable substrate for fermentation and methanogenesis once introduced O_2_ is consumed. Future studies of microbial growth and succession in subsurface permafrost would benefit from examining the impact of mixing, dispersal, oxygenation, and/or seeding from surface environments. Even in the absence of dispersal, our study demonstrates that resuscitated microorganisms have a remarkable capacity for growth and turnover, with the potential for dramatic reshaping of the microbial community during extended thaw conditions (**Fig 4**, **Fig. S9 - S13**).

A caveat of environmental molecular studies is the recalcitrance of target analytes (nucleic acids, lipids, proteins, etc.) following cell death (Carini et al., 2016). We target our study towards fatty acids derived from intact polar phospho- and glycolipids due to their rapid degradation upon cell death (Å. Frostegård et al., 2011; Lipp et al., 2008; Schubotz et al., 2009; Zhang et al., 2019). However, free PLFAs themselves could conceivably decay more slowly in low-temperature incubations due to hindered kinetics of IPL hydrolysis. By incubating over long durations (up to 6 months), our goal was to minimize this risk, as phospholipid headgroup cleavage occurs on the order of hours (Zhang et al., 2019). Of great concern is the recalcitrance of necromass-derived DNA in the environment, which has been demonstrated to account for a substantial fraction of DNA recovered for microbial community analysis (Carini et al., 2016). Thus, a fraction of DNA extracted in this study is likely representative of relict DNA contributing to 16S results. However, previous work at the Fox Permafrost Tunnel has shown that living microbial community composition is imprinted onto the relict DNA pool, implying that descriptions of microbial community structure at this site are not vastly different when relic DNA is removed (Burkert et al., 2019). Still, during microbial community succession, subtle changes to taxonomic composition of the community may be obscured by the persistence of relict DNA. Future studies to examine biomolecular persistence are needed to interpret genomic- and SIP-based studies more faithfully in cold environments.

One implication of this work is that the magnitude of ongoing warming appears less important than the lengthening of the thaw season for liberating recalcitrant C in subsurface permafost. Even with increasing warmth and a deeper active thaw layer, as long as the warm season is short enough (i.e., refreezing takes place before the resuscitation and growth lag time), subsurface permafrost is unlikely to be highly vulnerable to degradation. GHG in the short term may therefore be dominated by [ancient] gases entrained in the frozen matrix, rather than those freshly produced. Therefore, in addition to temperature, the seasonality and duration of the freeze/thaw interval may be a critical parameter to consider for predictions of biogeochemical responses to warming regions.

This study focuses on discontinuous, syngenetic surface and subsurface permafrost in interior Alaska. Many other kinds of permafrost exist in Alaska and elsewhere throughout the Arctic. For example, continuous permafrost on Alaska’s north slope and in Siberia extends hundreds of meters to over a kilometer into the subsurface. Whether permafrost at these extreme depths will exhibit similar rates of microbial resuscitation and growth given ongoing warming remains an open question. Furthermore, the limits of life in deep subsurface permafrost habitability have not been investigated. While some hyperarid soils have been described as essentially sterile, with microbial biosignatures falling below detection as conditions grow increasingly hostile (Dragone et al., 2021), it is unknown if similar gradients can be found in deep permafrost. This begs the question: in the deep subsurface permafrost, does there exist a fringe beyond which signatures of microbial activity can no longer be detected, and how will these deep horizons respond to a changing climate?

## 4 Conclusions

In this study, we quantify microbial growth rates and resuscitation in surface and subsurface permafrosts under a range of thaw scenarios. Across our samples, microbial resuscitation is detected as early as one week following thaw disturbance. However, the degree of microbial growth and proliferation varies substantially between surface and subsurface sites. Subsurface permafrosts exhibit extremely low rates of biomass turnover after 30 days and only 0.001 to 0.01% of the microbial community is resuscitated on a daily basis. In subsurface samples, production of glycolipids is more robust than phospholipids, and the opposite is true in surface samples, suggesting a key physiological role for glycolipids in long term cryoprotection during burial. Microbial resuscitation and growth does not clearly relate to the thaw temperature, indicating that factors other than temperature – such as nutrient availability – primarily control microbial resuscitation and biomass turnover via the community members primed to take advantage of local thaw conditions. In all thaw conditions (incubations at temperatures > 0°C), microbial communities undergo dramatic restructuring after 6 months, with previously trace microbial constituents becoming dominant members of the community. Finally, under the anaerobic conditions tested in this study, CO_2_ is emitted more rapidly in surface than subsurface permafrost. Conversely, CH_4_ is emitted more readily in subsurface permafrost. Taxonomic indicators of biological methanogenesis (i.e., the presence of *Methanobacterium* and other methanogenic archaea) do not clearly relate to the emission of CH_4_. Overall, gas emissions from subsurface permafrost samples do not follow clear trends with temperature. Our study is the first to apply stable hydrogen lipidomic stable isotope probing to permafrosts, and, to our knowledge, represents the first measurement of *in situ* microbial growth rates in subsurface permafrost. Future efforts deploying lipid-SIP in permafrost-hosted environments may further elucidate the timing of microbial resuscitation and growth under a wider range of environmental perturbations, and how anabolic activity maps onto ecosystem-scale carbon cycle processes and GHG emissions. This work represents an initial contribution to assessing biological activity in deep permafrost environments under warming scenarios, a task urgently required for assessing the biogeochemical implications of the ongoing warming of cold regions.

## Acknowledgements

We thank Noah Fierer and Kevin Rozmiarek for valuable discussions during the preparation of this manuscript. We thank Rachel Mackelprang, Joy O’Brien, Gareth Trubl, Karl Romanowicz, and Duane Froese for advice during the design of this study. Isotopoic determination of heavy water tracer solutions was conducted by the Feng Lab at Dartmouth College, with thanks to Wil Leavitt. Soil geochemistry was analyzed at the Colorado State University Soil and Water Testing Laboratory. Isotope ratio mass spectrometry was conducted at the CU Boulder Earth Systems Stable Isotope Lab (CUBES-SIL) core facility (RRID:SCR_019300). 16S amplification, cleanup, and sequencing was carried out at the University of Colorado Boulder Center for Microbial Exploration with thanks to Jessica Henley. T. Caro was supported by a NSF Graduate Research Fellowship. T. Caro and S. Jech were supported through the IQ Biology Program of the BioFrontiers Institute at the University of Colorado, Boulder. NOSAMS is supported by the NSF Cooperative Agreement number, OCE-1755125. This work was funded by a U.S. Army Research Office grant (W911NF2120119 / 78484-LS) awarded to S. Kopf. T. Douglas was supported by the Strategic Environmental Research and Development Program (project RC18-1170) and the Environmental Security Technology Demonstration Program (project RC22-7408). R. Barbato acknowledges funding from the US Department of Defense—PE 0602144A Program Increase ‘Defense Resiliency Platform Against Extreme Cold Weather’.

## Data Availability Statement

Raw data generated during this study is available at https://osf.io/fm5ux/. Processed data and data analysis scripts are available on Zenodo at https://doi.org/10.5281/zenodo.14624685.

