## Supplemental Information for "Microbial Resuscitation and Growth Rates in Deep Permafrost: Lipid Stable Isotope Probing Results from the Permafrost Research Tunnel in Fox, Alaska"

Tristan A. Caro[1]*, Ashley Maloney[1], Jamie M. McFarlin[2], Sierra Jech[3], Amanda Barker[4], Tom Douglas[4], Robyn Barbato[4], Sebastian H. Kopf[1]

1. Department of Geological Sciences, University of Colorado Boulder, Boulder CO, USA
2. Department of Geology and Geophysics, University of Wyoming, Laramie WY, USA
3. Department of Ecology and Evolutionary Biology, University of Colorado Boulder, Boulder CO, USA
4. US Army Cold Regions Research and Engineering Laboratory

*Corresponding Author
Tristan A. Caro

Benson Earth Sciences Bldg. Rm. 285, UCB 399
2200 Colorado Ave, Boulder, CO 80309

This file contains:

- Supplementary Text
- Supplementary Figures 1 - 13
- Supplementary Tables 1 - 3

### Supplementary Text

### Lipid Quantification

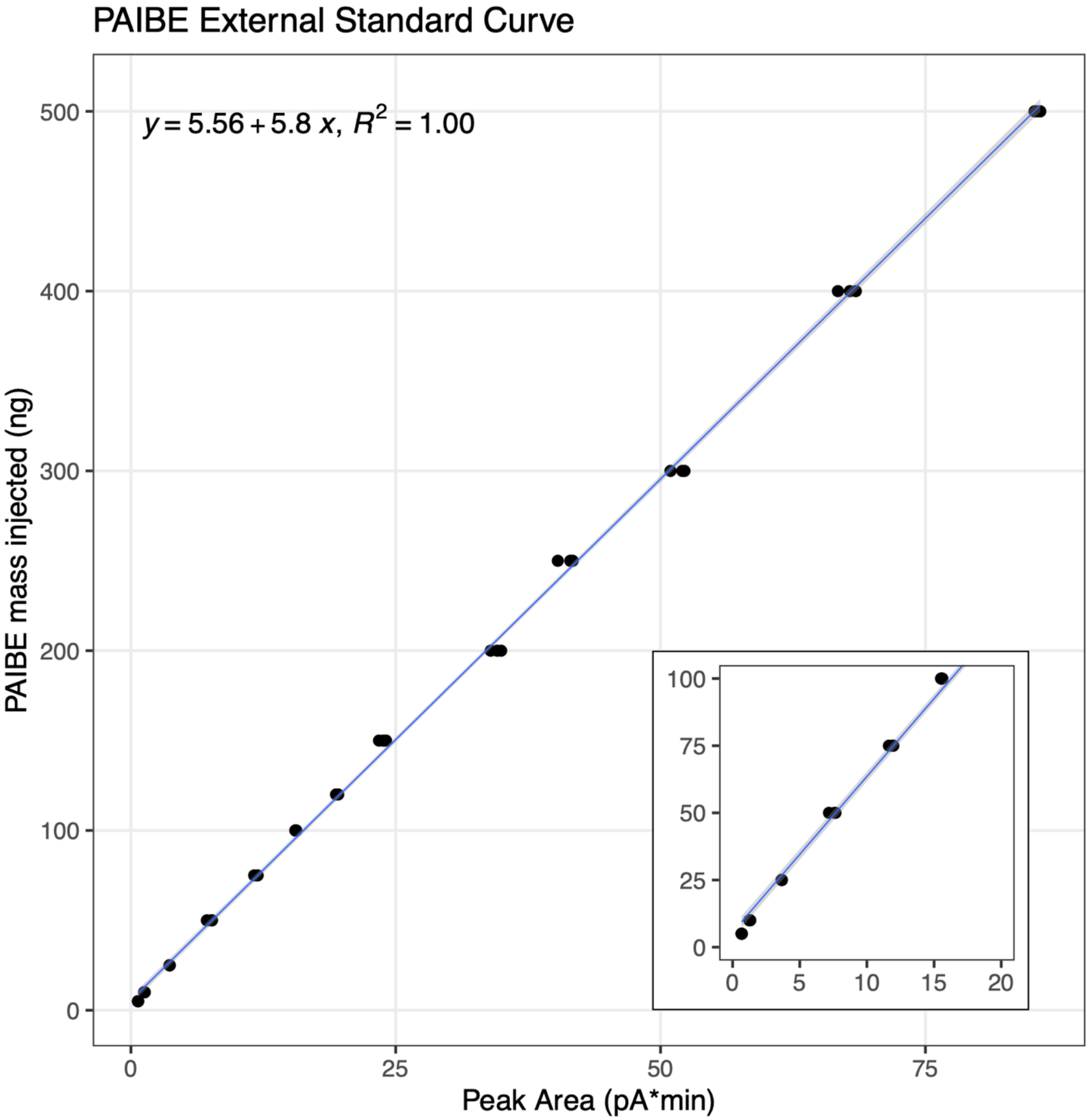


**Supplementary Figure S1.** Standard curve of palmitate isobutyl ester (PAIBE) analyzed by GC-FID. Inset panel displays a zoomed-in view of the bottom-left corner of the plot.

***Lipid quantification results***

Supplementary Figures S7 – S12 display results of lipid quantification by GC-FID. Note that for IRMS analyses, many of the GLFAs/PLFAs detected by GC-FID were not able to be quantified with acceptable error due to low signal intensities. Mono- and di-unsaturated FAMEs were aggregated due to uncertainty associated with identifying position of unsaturation. Supplementary

Figure S13 displays the external standard curve that was used for FAME quantification by GC-FID.

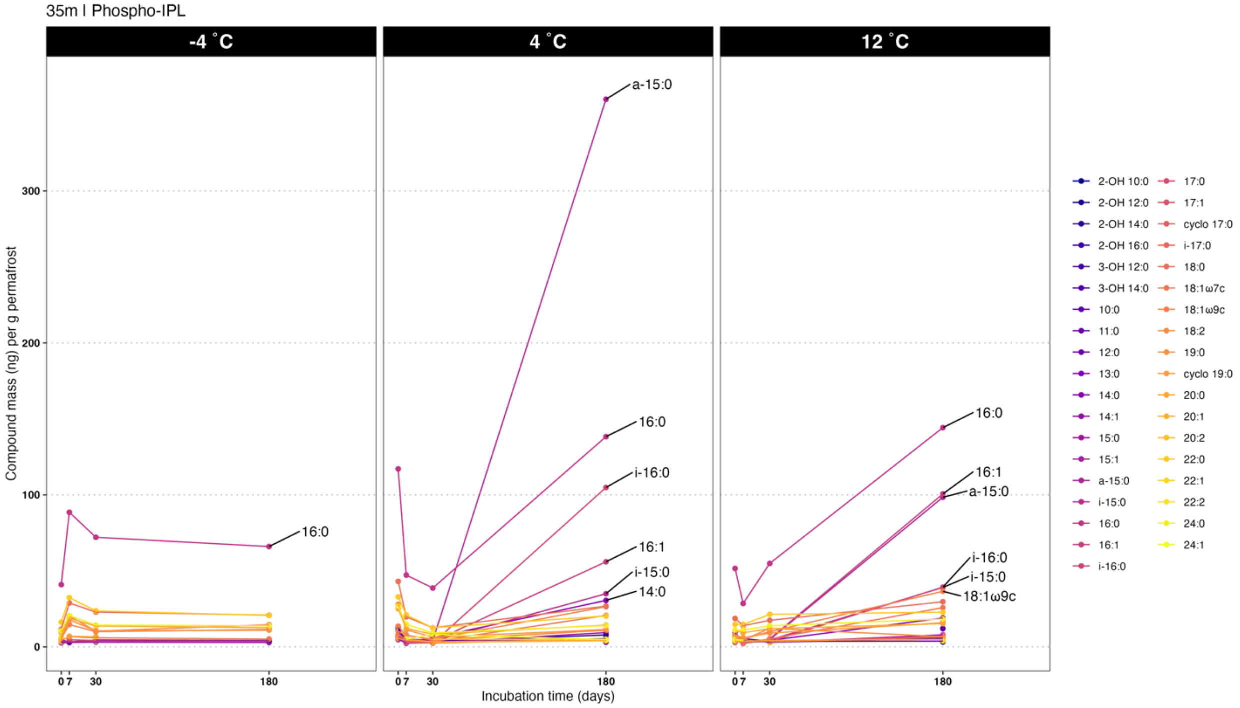


**Supplementary Figure S2.** Compound-specific masses of PLFA compounds in subsurface core 35m.

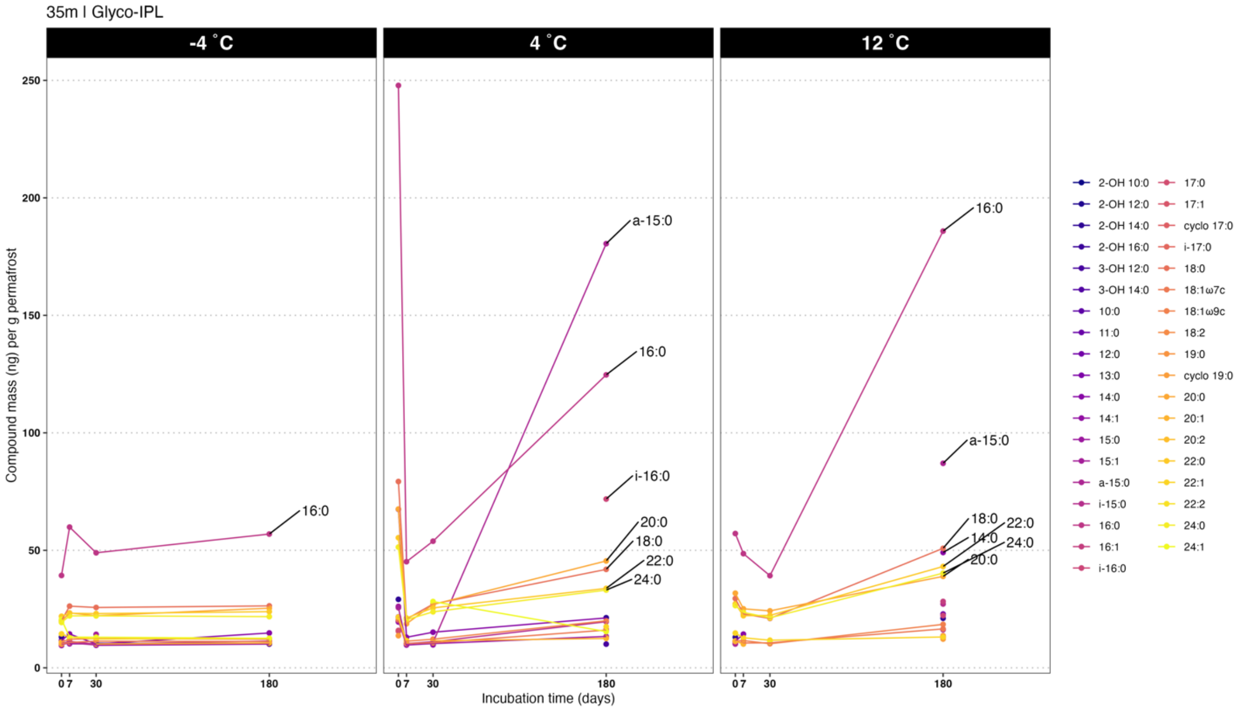


**Supplementary Figure S3.** Compound-specific masses of GLFA compounds in subsurface core 35m.

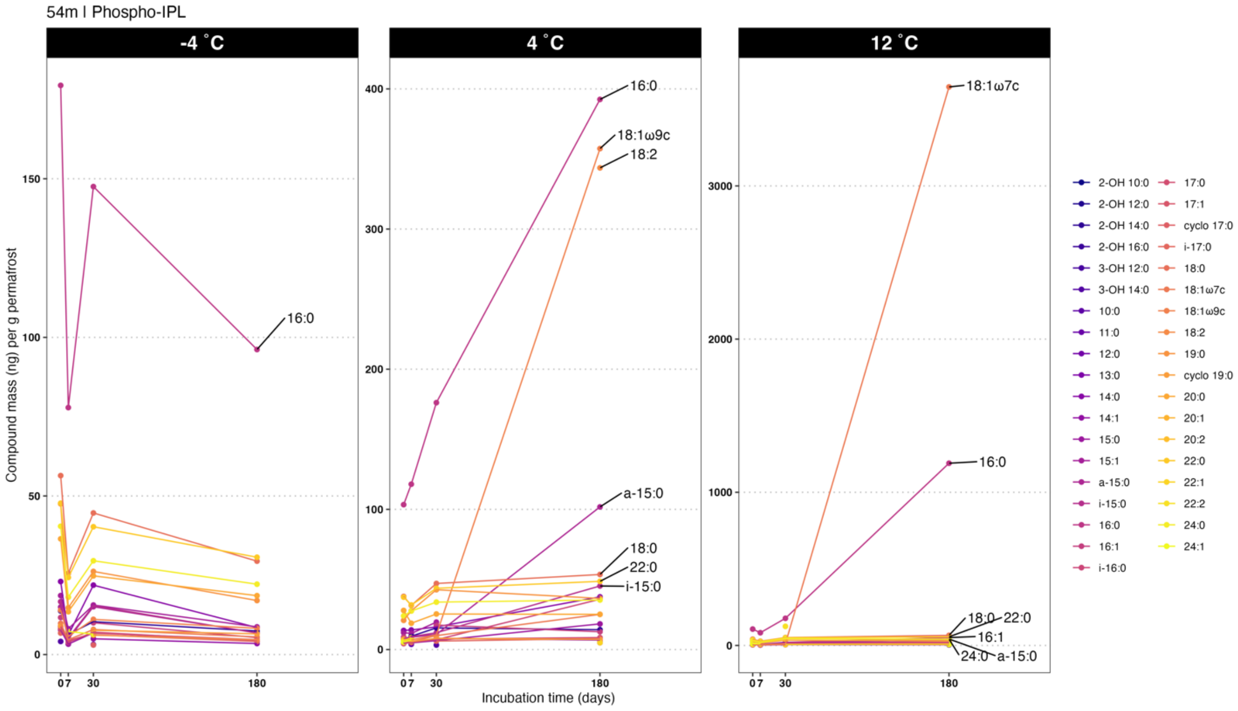


**Supplementary Figure S4.** Compound-specific masses of PLFA compounds in subsurface core 54m.

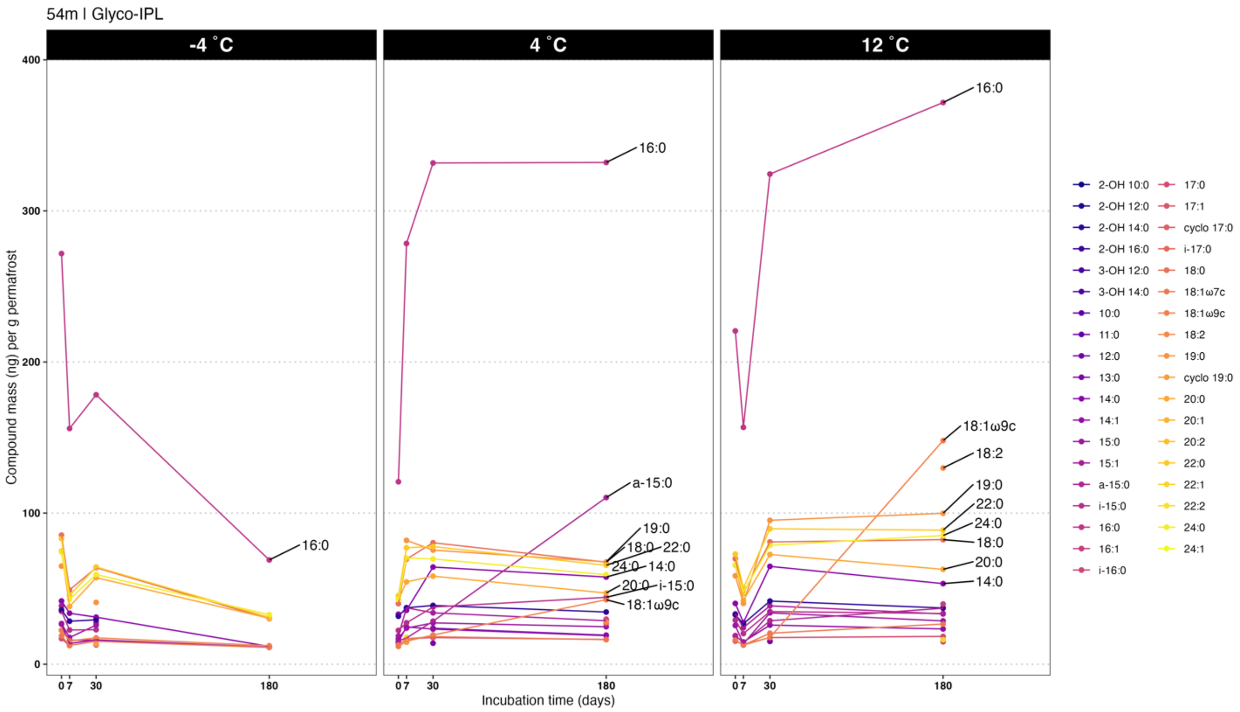


**Supplementary Figure S5.** Compound-specific masses of GLFA compounds in subsurface core 54m.

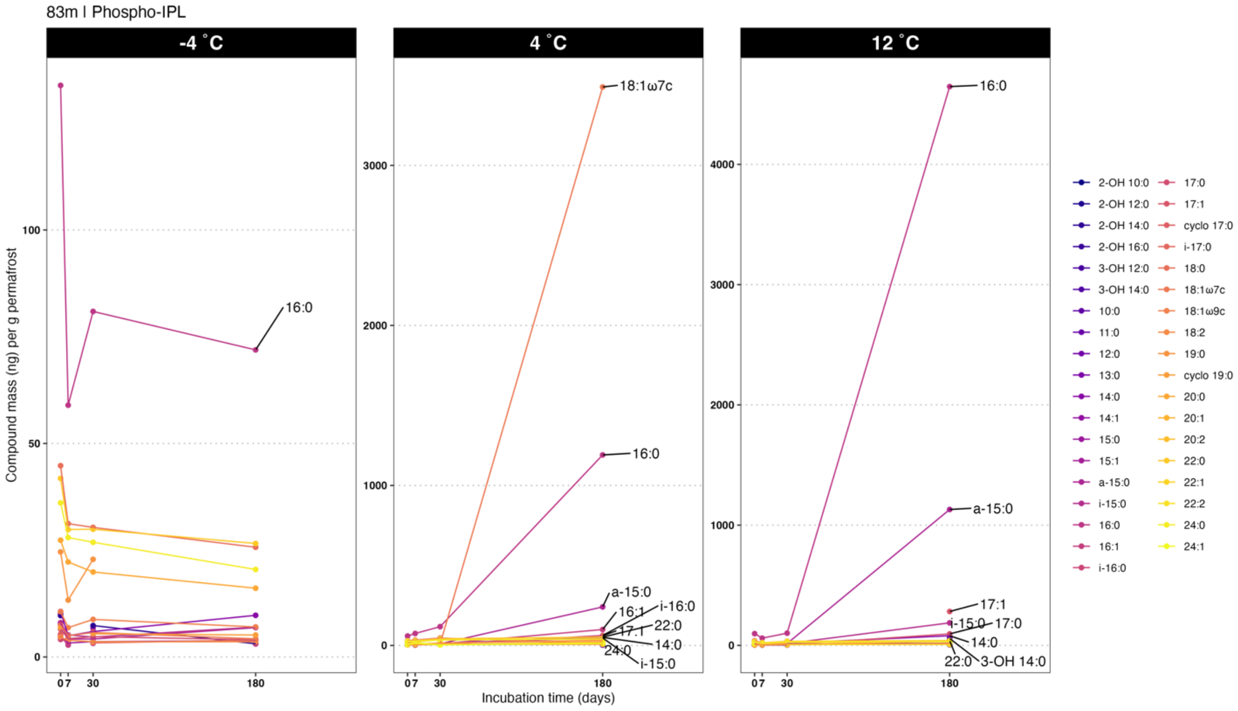


**Supplementary Figure S6.** Compound-specific masses of PLFA compounds in subsurface core 83m.

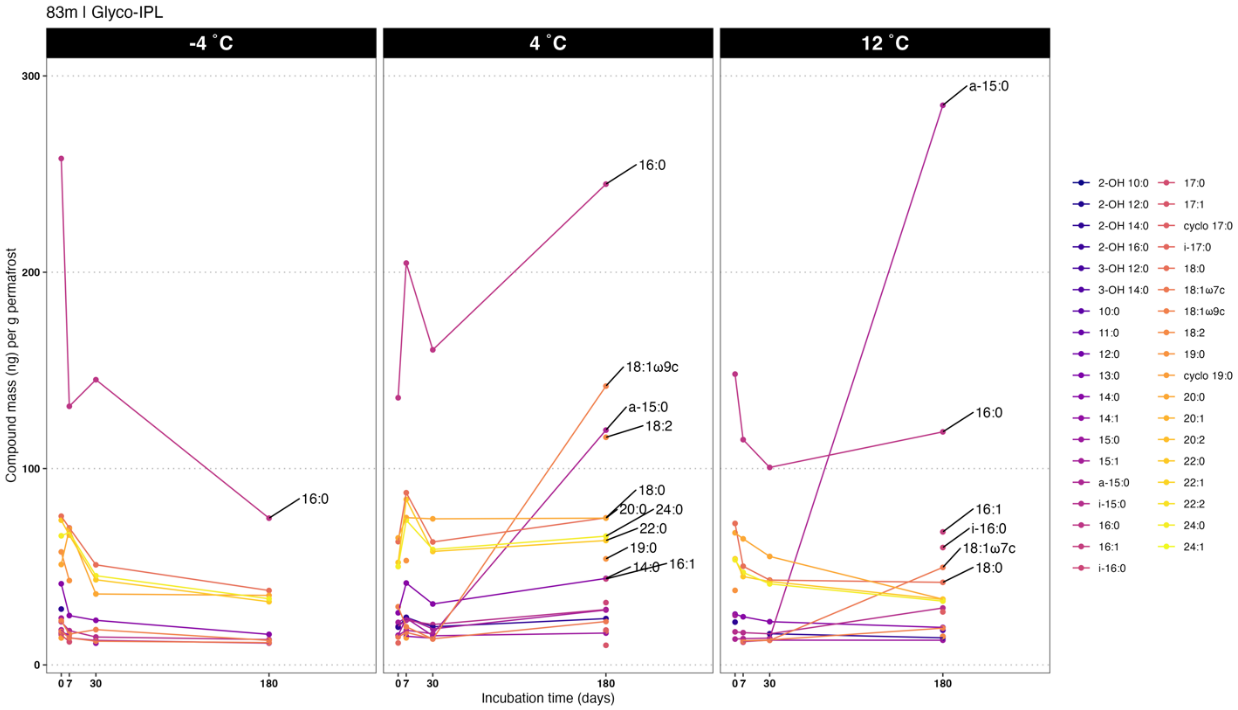


**Supplementary Figure S7.** Compound-specific masses of GLFA compounds in subsurface core 83m.

### 16S rRNA gene amplicon sequencing results

Figures S1 – S6 include display supplemental 16S data.

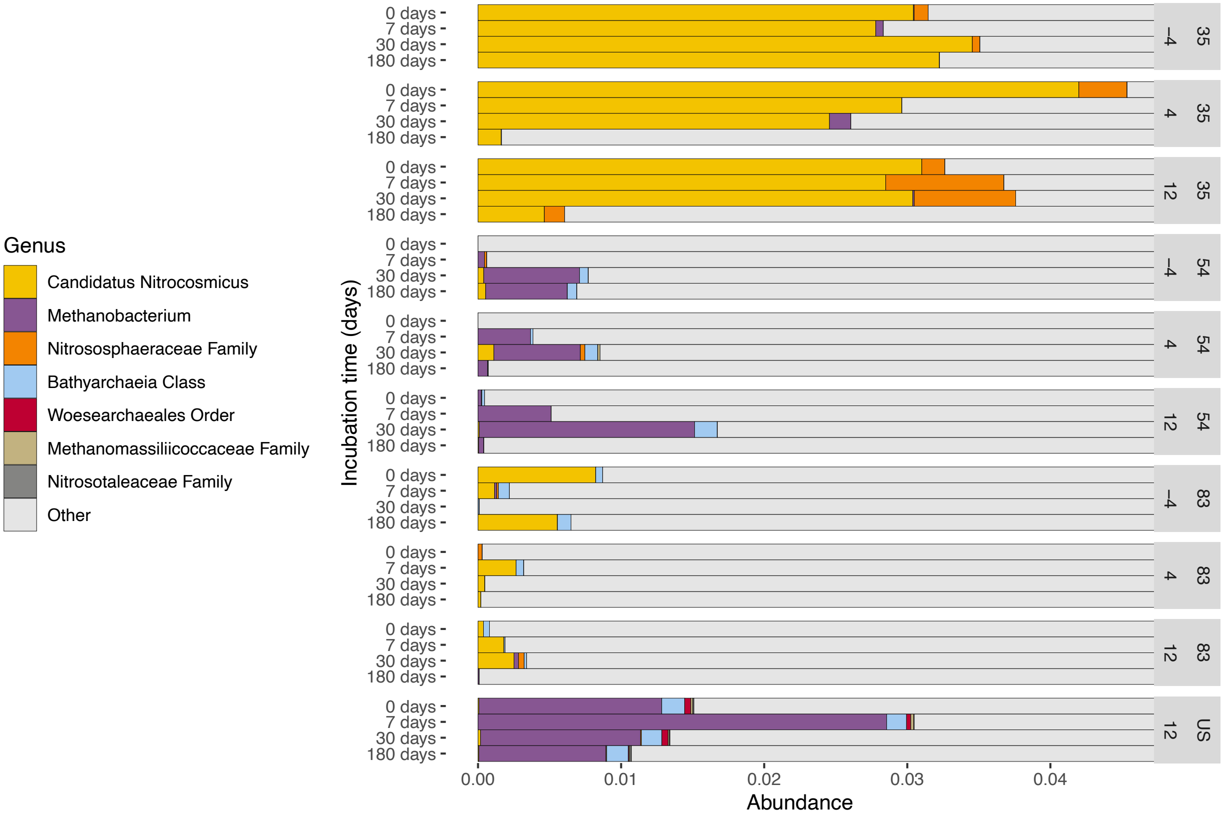


**Supplementary Figure S8.** 16S relative abundance of Archaea across samples. Horizontal axis indicates relative abundance estimated by 16S rRNA gene amplicon sequencing. “Other” indicates all taxa outside of domain Archaea.

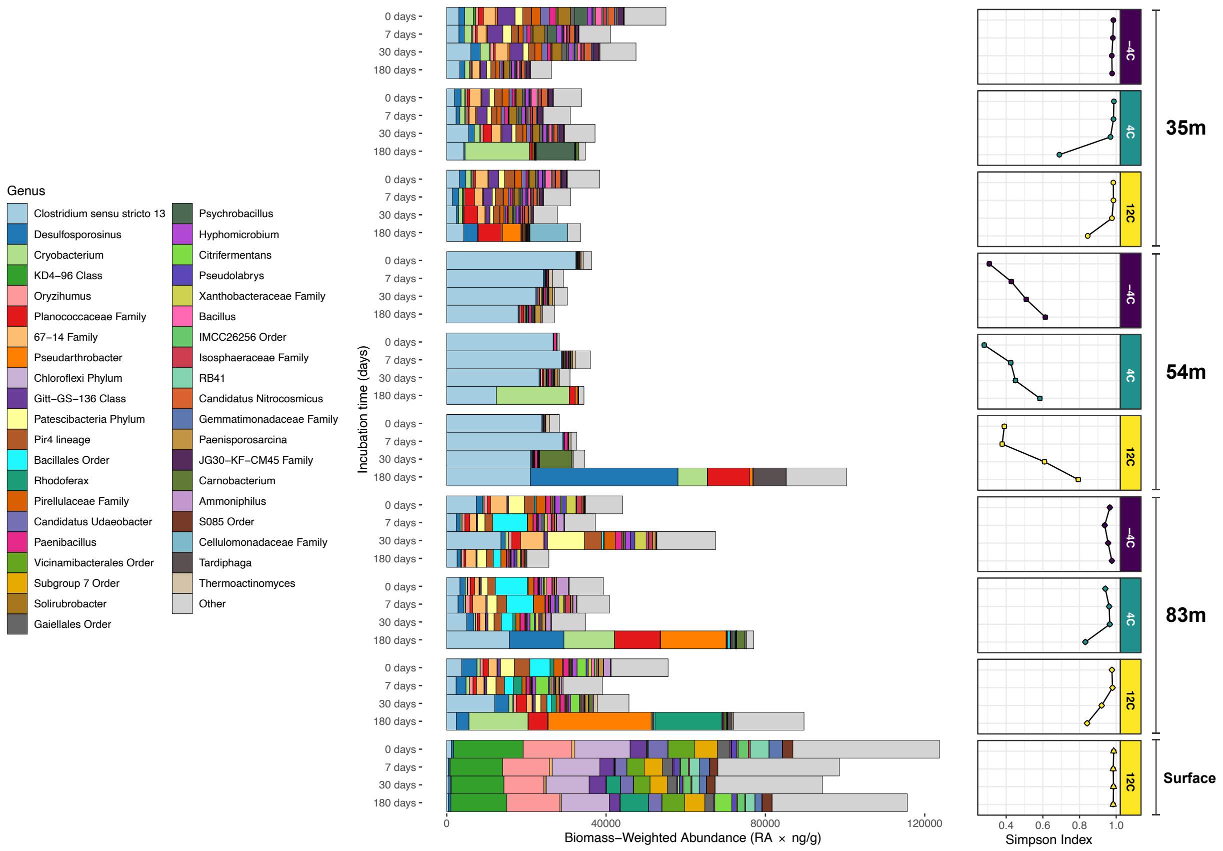


**Supplementary Figure S9.**

Barplot displays biomass-weighted abundances which are calculated as taxonomic relative abundance (RA), inferred by 16S rRNA gene amplicon sequencing, weighted by the total quantity of PLFA in the sample (ng PLFA per g permafrost). On the right is displayed the alpha (Simpson's) diversity index over the course of each incubation. Bar plot fill is by lowest annotated taxonomic rank down to Genus.

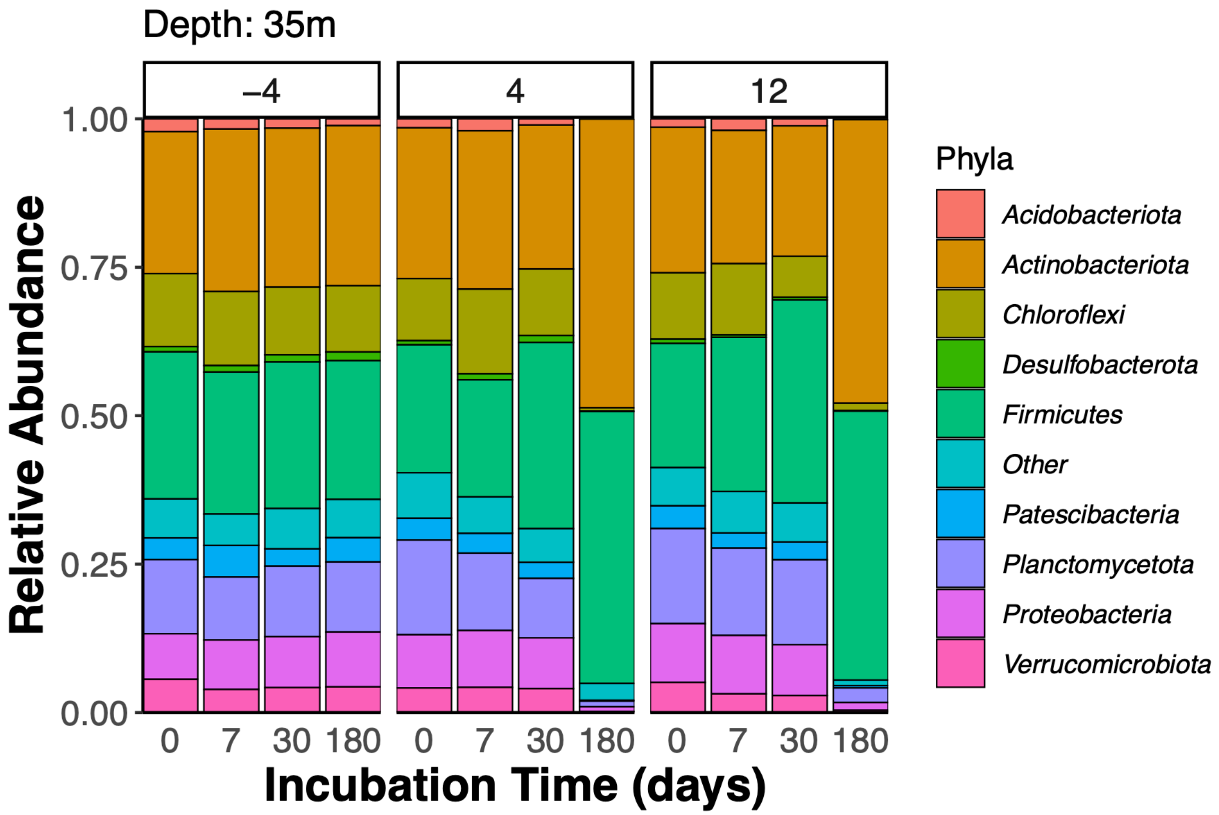


**Supplementary Figure S10. Relative abundance of microbial taxa in subsurface core 35m.** Taxa are grouped at the phylum level.

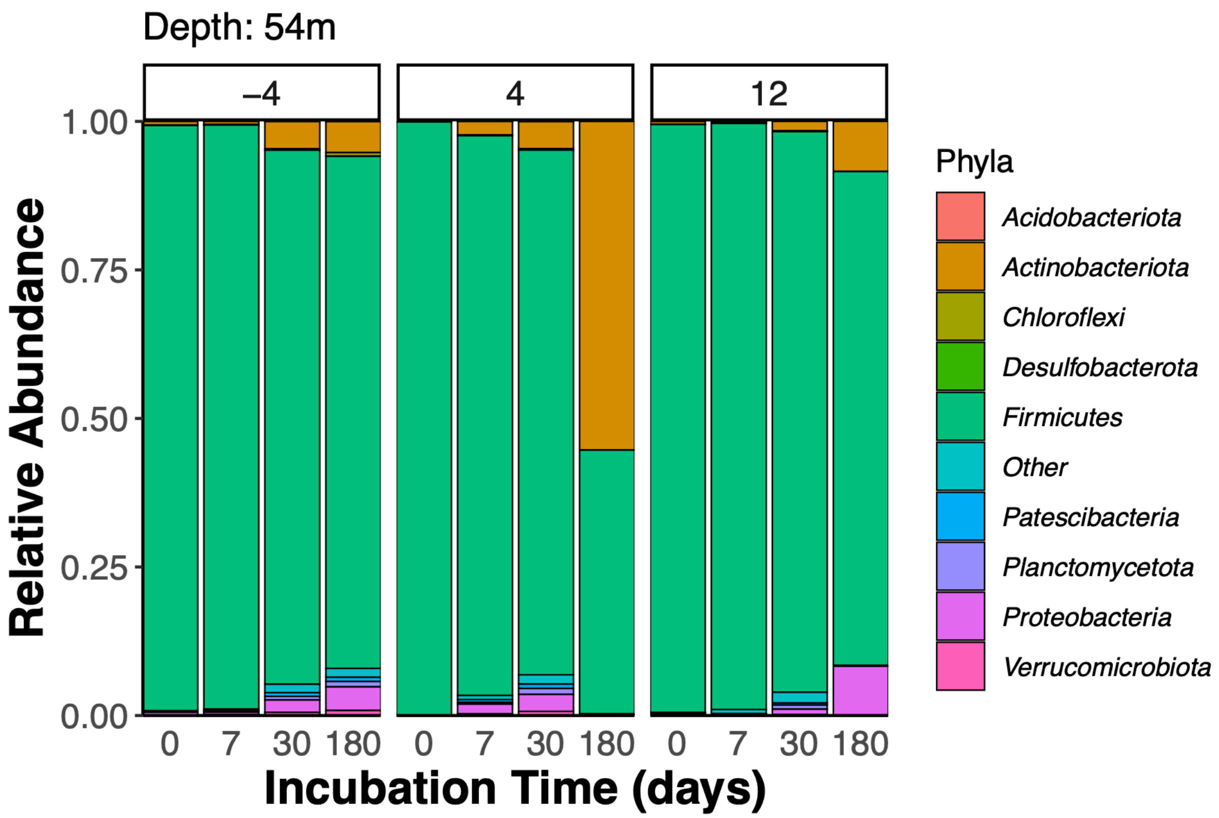


**Supplementary Figure S11. Relative abundance of microbial taxa in subsurface core 54m.** Taxa are grouped at the phylum level.

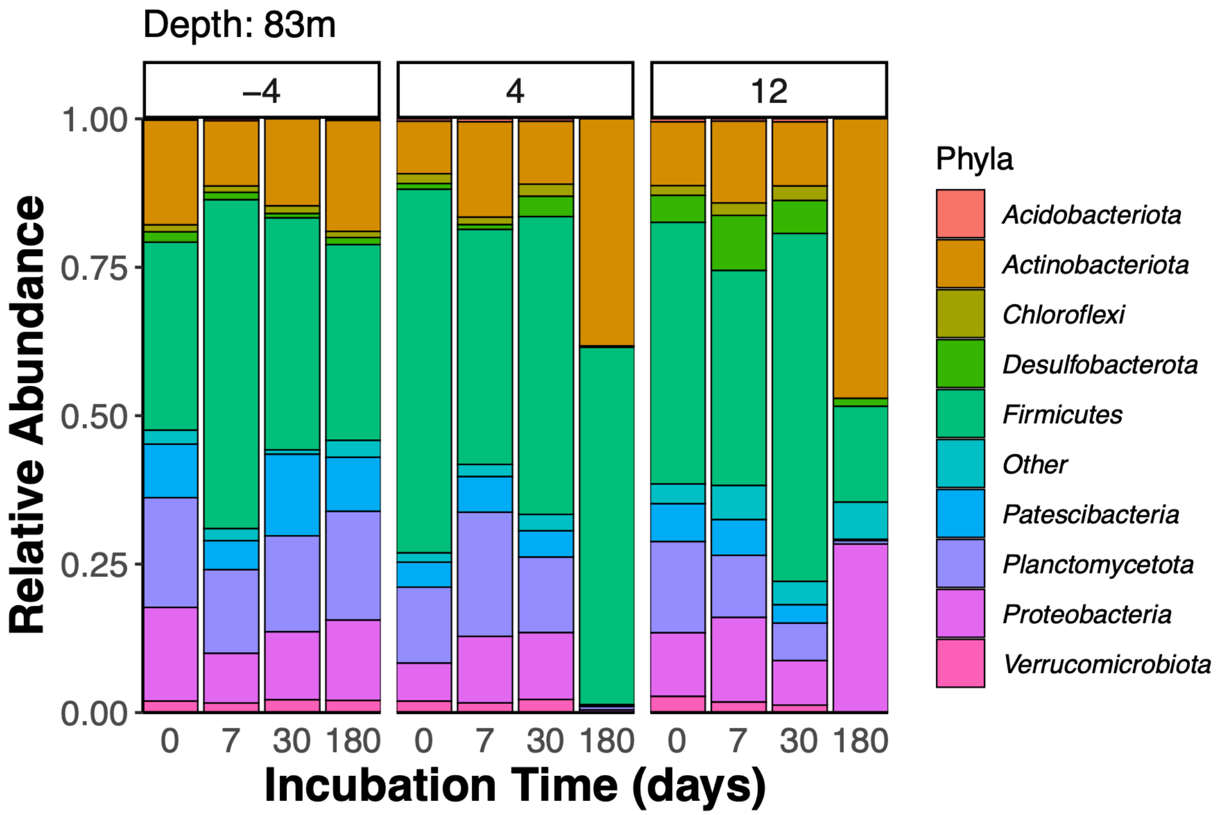


**Supplementary Figure S12. Relative abundance of microbial taxa in subsurface core 83m.** Taxa are grouped at the phylum level.

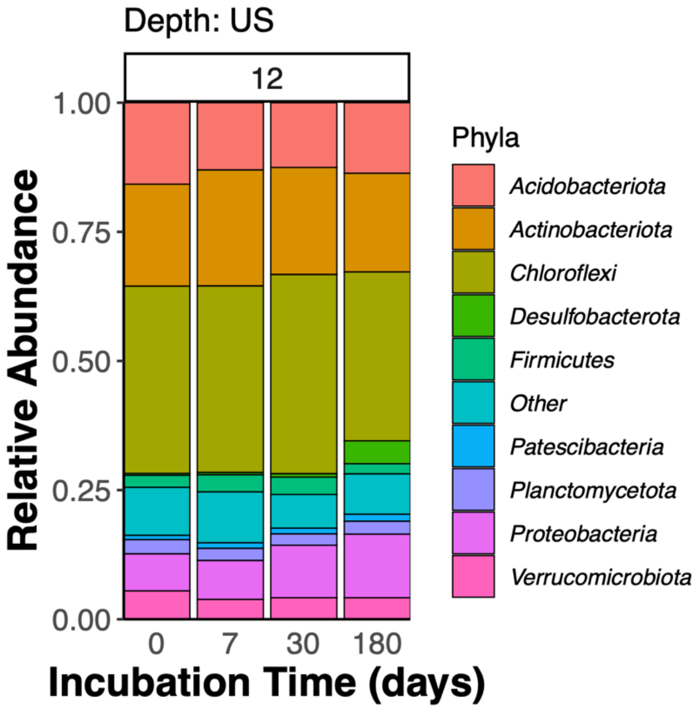


**Supplementary Figure S13. Relative abundance of microbial taxa in surface core "undisturbed surface" (US).** Site "US" is simply referred to as "surface permafrost core" in the main text. Taxa are grouped at the phylum level.

| Core | Incubation Temperature (C) | Incubation Time (days) | Growth Detected? | Growth Quantifiable? | Growth Rate ($days^{-1}$)[1] | GLFA Content (pg)[2] | GLFA turnover rate (pg/day)[1,2] | Cell density (cells/g)[2,3] | Cell turnover (cells/day)[1,3] |
|---|---|---|---|---|---|---|---|---|---|
| 35m | -4 | 7 | TRUE | TRUE | $2.3 \times 10^{-4}$ | $8.2 \times 10^{5}$ | 186.4 | $2.3 \times 10^{8}$ | $5.3 \times 10^{4}$ |
| 35m | 4 | 7 | TRUE | TRUE | $4.0 \times 10^{-4}$ | $1.4 \times 10^{6}$ | 560.8 | $4.0 \times 10^{8}$ | $1.6 \times 10^{5}$ |
| 35m | 4 | 30 | TRUE | TRUE | $5.1 \times 10^{-5}$ | $2.1 \times 10^{6}$ | 106.1 | $5.9 \times 10^{8}$ | $3.0 \times 10^{4}$ |
| 35m | 12 | 7 | TRUE | TRUE | $2.0 \times 10^{-4}$ | $1.1 \times 10^{6}$ | 216.5 | $3.2 \times 10^{8}$ | $6.2 \times 10^{4}$ |
| 35m | 12 | 30 | FALSE | FALSE | $2.4 \times 10^{-5}$ | $1.2 \times 10^{6}$ | 29.0 | $3.5 \times 10^{8}$ | $8.3 \times 10^{3}$ |
| 54m | -4 | 7 | TRUE | TRUE | $3.3 \times 10^{-4}$ | $1.3 \times 10^{6}$ | 448.1 | $3.8 \times 10^{8}$ | $1.3 \times 10^{5}$ |
| 54m | -4 | 30 | TRUE | TRUE | $6.3 \times 10^{-5}$ | $1.2 \times 10^{6}$ | 72.6 | $3.3 \times 10^{8}$ | $2.1 \times 10^{4}$ |
| 54m | 4 | 7 | TRUE | TRUE | $1.7 \times 10^{-4}$ | $1.9 \times 10^{6}$ | 320.4 | $5.4 \times 10^{8}$ | $9.1 \times 10^{4}$ |
| 54m | 4 | 30 | TRUE | TRUE | $6.7 \times 10^{-5}$ | $1.2 \times 10^{6}$ | 82.3 | $3.5 \times 10^{8}$ | $2.3 \times 10^{4}$ |
| 54m | 12 | 7 | TRUE | TRUE | $3.0 \times 10^{-4}$ | $1.7 \times 10^{6}$ | 502.9 | $4.8 \times 10^{8}$ | $1.4 \times 10^{5}$ |
| 54m | 12 | 30 | FALSE | FALSE | $7.4 \times 10^{-5}$ | $1.4 \times 10^{6}$ | 106.0 | $4.1 \times 10^{8}$ | $3.0 \times 10^{4}$ |
| 83m | -4 | 7 | TRUE | TRUE | $1.9 \times 10^{-4}$ | $3.0 \times 10^{6}$ | 561.5 | $8.5 \times 10^{8}$ | $1.6 \times 10^{5}$ |
| 83m | -4 | 30 | TRUE | TRUE | $5.3 \times 10^{-5}$ | $4.1 \times 10^{6}$ | 218.9 | $1.2 \times 10^{9}$ | $6.2 \times 10^{4}$ |
| 83m | 4 | 7 | TRUE | TRUE | $2.3 \times 10^{-4}$ | $2.7 \times 10^{6}$ | 624.8 | $7.6 \times 10^{8}$ | $1.8 \times 10^{5}$ |
| 83m | 4 | 30 | TRUE | TRUE | $5.0 \times 10^{-5}$ | $1.4 \times 10^{6}$ | 72.2 | $4.1 \times 10^{8}$ | $2.1 \times 10^{4}$ |
| 83m | 12 | 7 | TRUE | TRUE | $2.6 \times 10^{-4}$ | $2.1 \times 10^{6}$ | 533.8 | $5.9 \times 10^{8}$ | $1.5 \times 10^{5}$ |
| 83m | 12 | 30 | TRUE | TRUE | $4.4 \times 10^{-5}$ | $1.5 \times 10^{6}$ | 68.2 | $4.4 \times 10^{8}$ | $1.9 \times 10^{4}$ |

[1] Values exhibit high error if 'Growth Quantifiable' is FALSE, take values as approximations.

[2] Reported per gram of permafrost.

[3] Estimated using PLFA/cell conversion factors reported in reference #.

**Supplementary Table S1. Glycolipid fatty acid turnover in subsurface cores.**

| Core | Incubation Temperature (C) | Incubation Time (days) | Growth Detected? | Growth Quantifiable? | Growth Rate (days −1)[1] | PLFA Content (pg)[2] | PLFA turnover rate (pg/day)[1,2] | Cell density (cells/g)[2,3] | Cell turnover (cells/day)[1,3] |
|---|---|---|---|---|---|---|---|---|---|
| 35m | -4 | 7 | TRUE | TRUE | $1.3 \times 10^{-4}$ | $4.0 \times 10^{5}$ | 51.8 | $1.1 \times 10^{8}$ | $1.5 \times 10^{4}$ |
| 35m | -4 | 30 | TRUE | TRUE | $3.3 \times 10^{-5}$ | $8.2 \times 10^{5}$ | 27.5 | $2.3 \times 10^{8}$ | $7.8 \times 10^{3}$ |
| 35m | 4 | 30 | TRUE | TRUE | $7.5 \times 10^{-5}$ | $6.8 \times 10^{5}$ | 50.5 | $1.9 \times 10^{8}$ | $1.4 \times 10^{4}$ |
| 35m | 12 | 30 | FALSE | FALSE | $7.9 \times 10^{-5}$ | $4.0 \times 10^{5}$ | 31.5 | $1.1 \times 10^{8}$ | $9.0 \times 10^{3}$ |
| 54m | -4 | 7 | FALSE | FALSE | $9.8 \times 10^{-5}$ | $3.3 \times 10^{5}$ | 32.0 | $9.3 \times 10^{7}$ | $9.1 \times 10^{3}$ |
| 54m | -4 | 30 | FALSE | FALSE | $3.4 \times 10^{-7}$ | $4.7 \times 10^{5}$ | 0.2 | $1.3 \times 10^{8}$ | $4.6 \times 10^{1}$ |
| 54m | 4 | 7 | TRUE | TRUE | $1.2 \times 10^{-4}$ | $5.3 \times 10^{5}$ | 62.2 | $1.5 \times 10^{8}$ | $1.8 \times 10^{4}$ |
| 54m | 4 | 30 | FALSE | FALSE | $4.7 \times 10^{-5}$ | $5.3 \times 10^{5}$ | 25.1 | $1.5 \times 10^{8}$ | $7.2 \times 10^{3}$ |
| 54m | 12 | 7 | FALSE | FALSE | $5.4 \times 10^{-5}$ | $4.3 \times 10^{5}$ | 22.9 | $1.2 \times 10^{8}$ | $6.5 \times 10^{3}$ |
| 54m | 12 | 30 | TRUE | TRUE | $6.6 \times 10^{-5}$ | $5.4 \times 10^{5}$ | 35.5 | $1.5 \times 10^{8}$ | $1.0 \times 10^{4}$ |
| 83m | -4 | 7 | FALSE | FALSE | 0.0 | $6.6 \times 10^{5}$ | 0.0 | $1.9 \times 10^{8}$ | 0.0 |
| 83m | -4 | 30 | FALSE | FALSE | 0.0 | $1.5 \times 10^{6}$ | 0.0 | $4.2 \times 10^{8}$ | 0.0 |
| 83m | 4 | 7 | FALSE | FALSE | 0.0 | $1.5 \times 10^{6}$ | 0.0 | $4.4 \times 10^{8}$ | 0.0 |
| 83m | 4 | 30 | FALSE | FALSE | 0.0 | $5.6 \times 10^{5}$ | 0.0 | $1.6 \times 10^{8}$ | 0.0 |
| 83m | 12 | 7 | FALSE | FALSE | $4.0 \times 10^{-5}$ | $6.5 \times 10^{5}$ | 25.7 | $1.9 \times 10^{8}$ | $7.3 \times 10^{3}$ |
| 83m | 12 | 30 | TRUE | TRUE | $5.7 \times 10^{-4}$ | $7.5 \times 10^{5}$ | 427.2 | $2.1 \times 10^{8}$ | $1.2 \times 10^{5}$ |

[1] Values exhibit high error if 'Growth Quantifiable' is FALSE, take values as approximations.

[2] Reported per gram of permafrost.

[3] Estimated using PLFA/cell conversion factors reported in reference #.

**Supplementary Table S2. Phospholipid fatty acid turnover in subsurface cores.**

| Core | Lipid Class | Incubation Time (days) | Incubation Temperature (C) | Growth Detected? | Growth Quantifiable? | Growth Rate (days −1)[1] | LFA Content (pg)[2] | PLFA turnover rate (pg/day)[1,2] | Cell density (cells/g)[2,3] | Cell turnover (cells/day)[1,3] |
|---|---|---|---|---|---|---|---|---|---|---|
| Surface | Glyco-IPL | 7 | 12 | FALSE | FALSE | $4.1 \times 10^{-4}$ | $6.0 \times 10^{6}$ | $2.5 \times 10^{3}$ | $1.7 \times 10^{9}$ | $7.1 \times 10^{5}$ |
| Surface | Phospho-IPL | 7 | 12 | TRUE | TRUE | $2.1 \times 10^{-3}$ | $5.3 \times 10^{6}$ | $1.1 \times 10^{4}$ | $1.5 \times 10^{9}$ | $3.1 \times 10^{6}$ |
| Surface | Glyco-IPL | 30 | 12 | TRUE | TRUE | $3.7 \times 10^{-4}$ | $3.8 \times 10^{6}$ | $1.4 \times 10^{3}$ | $1.1 \times 10^{9}$ | $4.0 \times 10^{5}$ |
| Surface | Phospho-IPL | 30 | 12 | TRUE | TRUE | $7.6 \times 10^{-3}$ | $7.1 \times 10^{6}$ | $5.4 \times 10^{4}$ | $2.0 \times 10^{9}$ | $1.6 \times 10^{7}$ |

[1] Values exhibit high error if 'Growth Quantifiable' is FALSE, take values as approximations.

[2] Reported per gram of permafrost.

[3] Estimated using PLFA/cell conversion factors reported in reference #.

**Supplementary Table S3. Lipid turnover in surface core.**
